# Serotonergic anxiety is a stress intensity-dependent state mediated by dopaminergic signaling

**DOI:** 10.64898/2026.08.28.747812

**Authors:** Zoltan K Varga, Jorun Van Mieghem, Lucia Jimenez-Fernandez, Adam Ujhelyi, Florence Kermen

**Affiliations:** Department of Neuroscience, Faculty of Health and Medical Sciences, University of Copenhagen, Copenhagen, Denmark

## Abstract

Despite the central role attributed to serotonin in anxiety, its involvement and therapeutic efficacy are inconsistent, raising the possibility that serotonergic recruitment depends on the stress history from which anxiety emerges. Using larval zebrafish, we combined graded glucocorticoid exposure with chemogenetic DRN manipulation, whole-brain activity mapping, and pharmacology to test whether stress intensity determines serotonergic involvement in anxiety.

Increasing stress intensity did not simply increase anxiety severity but generated distinct anxiety phenotypes. Lower glucocorticoid exposure produced context-general anxiety that required the serotonergic DRN, whereas higher exposure produced context-dependent, DRN-independent anxiety. Whole-brain mapping identified a posterior tubercular/hypothalamic dopaminergic region associated with DRN-dependent anxiolysis, while D1, but not D2 receptor antagonism abolished the anxiolytic effect of DRN ablation.

Together, our findings identify stress intensity as a determinant of serotonergic recruitment during anxiety, reveal a downstream contribution of D1-dependent dopaminergic signaling, and provide a framework for understanding the mechanistic heterogeneity of anxiety and its variable response to serotonergic treatments.

**GRAPHICAL ABSTRACT:** 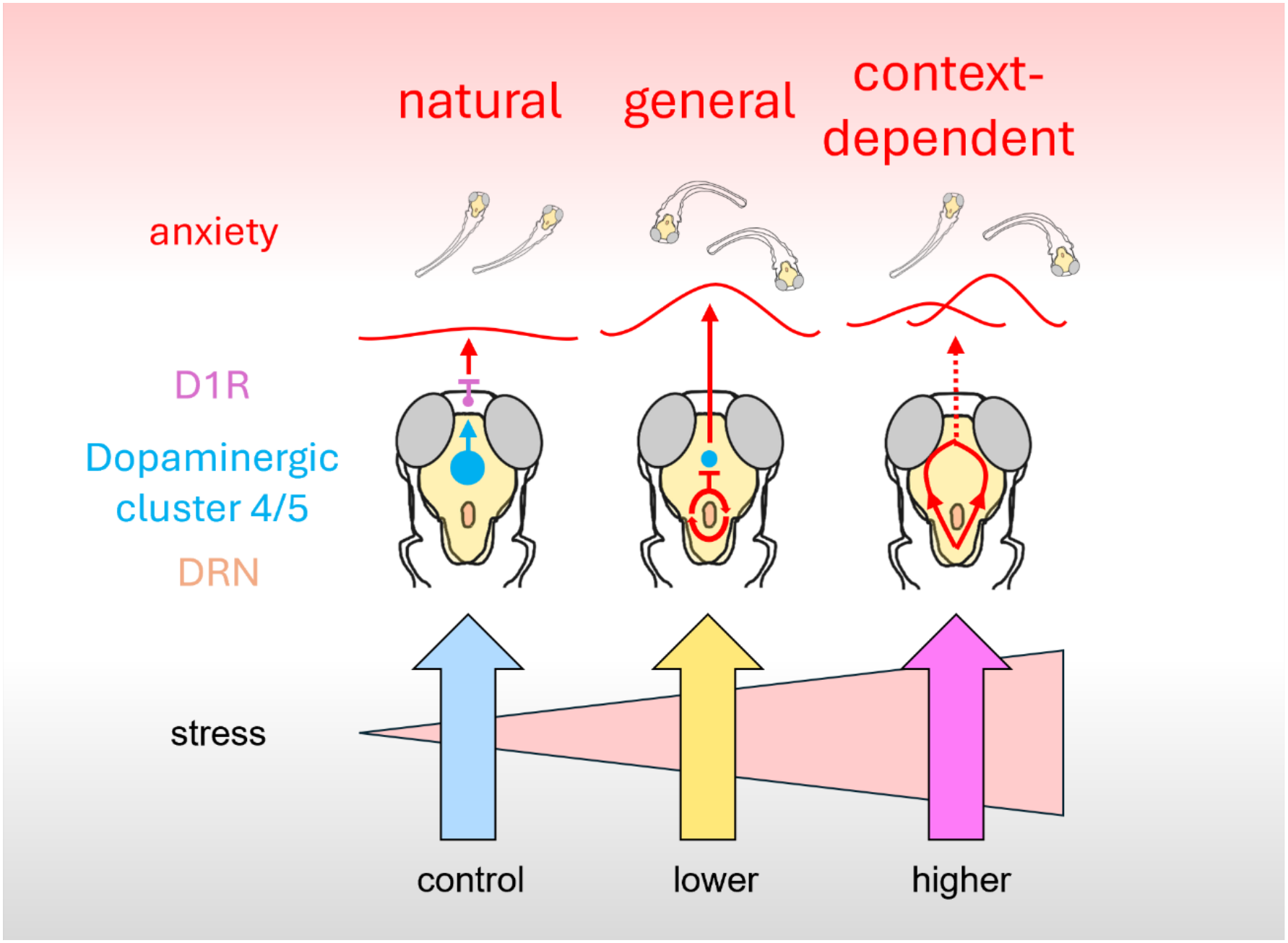

## INTRODUCTION

Anxiety disorders are one of the fastest-growing global health challenges^1^ and account for nearly half of all mental disorders^2,3^. Despite their high prevalence, the neurobiological mechanisms underlying anxiety remain incompletely understood, and treatment responses are highly variable^4^.

Both the mechanistic understanding and pharmacological treatment of anxiety have largely centered on serotonergic signaling^5,6^. Although extensive pharmacological and genetic evidence supports a role for serotonin in anxiety regulation^7,8^, numerous studies report weak, absent, or even paradoxical serotonergic effects^9–14^, and serotonergic anti-anxiety treatments often show variable efficacy^15,16^. One proposed explanation is that only specific components of the serotonergic system contribute to anxiety^9,10,17,18^. While this may partly account for the observed inconsistencies, it does not address a more fundamental question: does serotonergic signaling contribute to all forms of anxiety, or only to specific anxiety states?

In addition to genetic factors, prior stress experiences are major determinants of anxiety and its treatment response. Increasing evidence suggests that stress history can modify serotonergic involvement in anxiety. For example, previous social instability or social isolation in rats, mice or zebrafish can enhance the anxiolytic efficacy of buspirone and SSRIs^19–23^, while previous neutral experience can decrease it. Conversely, chronic unpredictable stress (CUS) induces an anxiety-like state in zebrafish that is independent of the dorsal raphe nucleus (DRN)^11^, the principal source of serotonin in vertebrates. Together, these findings suggest that prior stress experiences might influence whether serotonergic mechanisms are involved in anxiety. The modality (social vs non-social)^17,24^ and duration^10^ of stress have both been proposed as key determinants of serotonergic involvement. However, serotonergic signaling also contributes to anxiety following non-social stress, such as restraint, or modulates anxiety under non-social, novelty challenges^22^, suggesting a more general mechanism. We therefore hypothesized that stress intensity determines the recruitment of serotonergic mechanisms during anxiety. Although such a framework could explain substantial variability in both anxiety pathophysiology and treatment responses, it has not been systematically tested experimentally.

Larval zebrafish (*Danio rerio*) is a powerful vertebrate model for testing this hypothesis. They exhibit conserved stress responses^25^, serotonergic circuitry^26,27^, and sensitivity to clinically relevant anxiolytic drugs^28–30^ while enabling high-throughput behavioral phenotyping^28^ combined with whole-brain mapping of neural activity^31,32^. These features allow systematic comparison of the neural mechanisms underlying anxiety states induced by different stress experiences.

Here, we investigated whether stress intensity determines the recruitment of serotonergic mechanisms during anxiety. Using graded glucocorticoid exposure, chemogenetic manipulation of the dorsal raphe nucleus (DRN), and whole-brain activity mapping in larval zebrafish, we identified a stress intensity-dependent serotonergic anxiety state and investigated downstream underlying mechanisms. We found that exposure to lower and higher hydrocortisone concentrations generated distinct anxiety-like phenotypes, with only the former depending on the DRN, as demonstrated by phenotypic rescue via DRN ablation. Rescue of this DRN-dependent anxiety state was associated with increased activity of hypothalamic/posterior tubercular dopaminergic neurons and required dopamine D1 receptor signaling. Together, our findings indicate that the involvement of serotonergic signaling in anxiety depends on the intensity of previous stress experience and identify a downstream dopaminergic pathway underlying this form of anxiety. These results provide a potential mechanistic framework for understanding variability in serotonergic treatment responses.

## RESULTS

### Experiment 1: Developmental exposure to the glucocorticoid hydrocortisone induces concentration-dependent anxiety

To establish a high-throughput model of stress-induced anxiety, we investigated the effects of hydrocortisone (HC) exposure at different concentrations and durations using the swimming plus-maze (SPM) test. Subjects were exposed to 0 (control), 0.5 (lower intensity stress), or 1 μM (higher intensity stress) HC in a water bath twice daily for 20 min between 6 and 12 dpf (Figure 1A–B). Behavioral assessments were performed on test-naive animals before treatment onset (day 1) and after 1, 4, and 7 days of exposure (days 2, 5, and 8, respectively).

**Figure 1.**
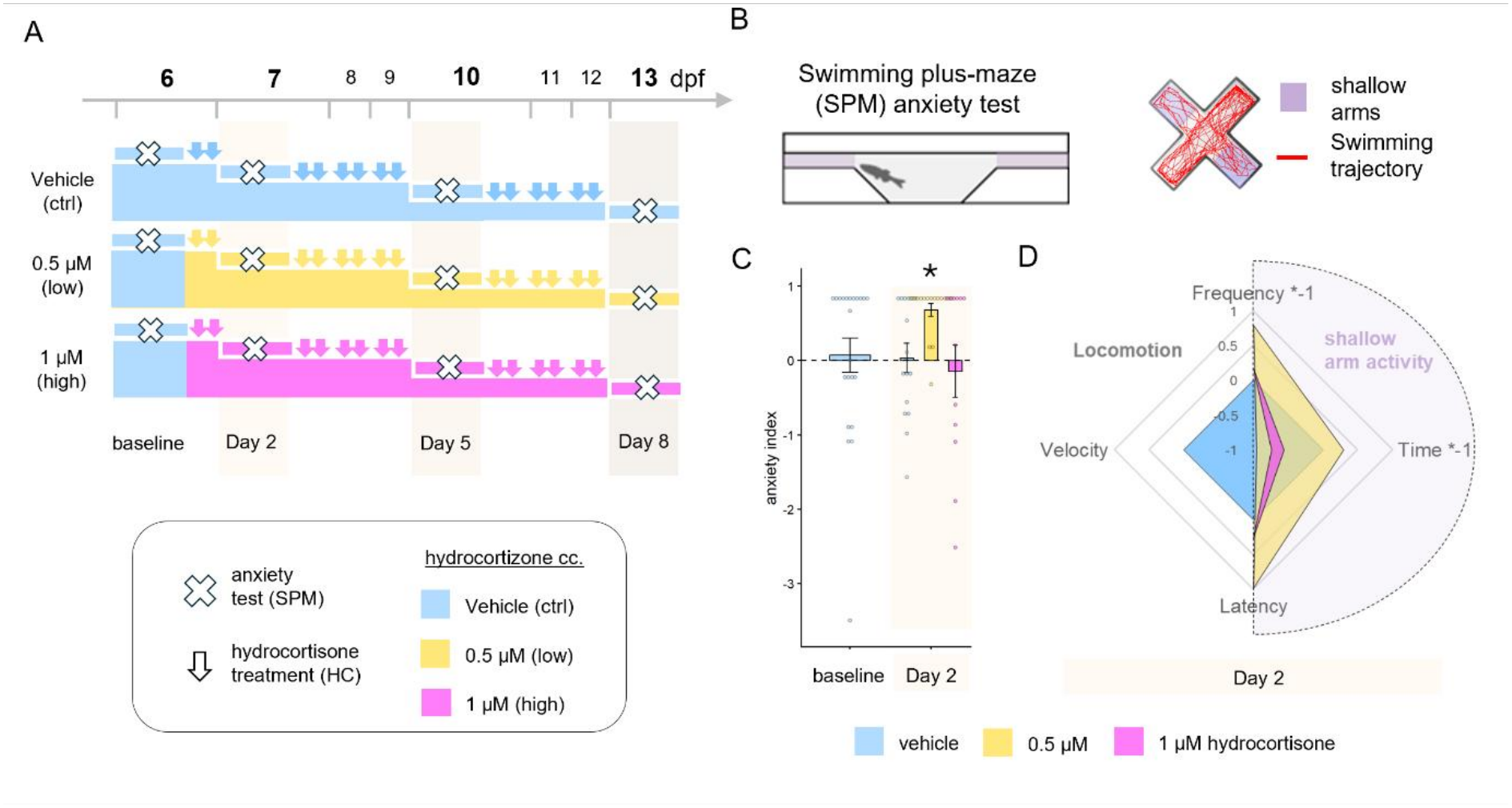
Concentration-and duration-dependent effects of hydrocortisone stress on anxiety. **A)** Treatment and sampling design of Experiment 1. **B)** Top and intersection view of the swimming plus-maze anxiety test. **C)** Anxiety score ((shallow arm entries *-1 + shallow arm time *-1 + shallow arm latency)/3) that positively reflects anxiety before and after treatments of hydrocortisone. **D)** Radial plot of the behavioral phenotype. Mean velocity and variables of shallow arm activity (source variables of the anxiety score) at day 2. Note that shallow arm entries (frequency) and time is multiplied by −1 so that all shallow arm variables positively reflects anxiety. Variables are scaled and normed to the ctrl group. n=16 fish per SPM sampling point.

The anxiety index – derived from time spent in, frequency of entries into, and latency to enter the shallow arms – was significantly increased in the lower-HC group on day 2 and returned to baseline thereafter (Figure 1C–D, Supplementary Figure 1D, Table 1). Individual SPM measures are shown in Supplementary Figure 1 and Supplementary table 1. Anxiety remained stable across sampling days in control animals (Supplementary Figure 1D). In contrast, locomotor activity was transiently reduced in control animals on day 2 relative to baseline and recovered at later time points (Supplementary Figure 1E). Both HC treatments further suppressed locomotion on day 2, with the effect gradually diminishing over subsequent assessments.

**Table 1:** Statistical summary of Experiment 1.

| **Exp.** | **Dependent variable** | **Analysis / Effect** | **df** | **Statistic** | ***P*** |
|---|---|---|---|---|---|
| **1** | **Anxiety index** | **Two-way ANOVA** |  |  |  |
|  |  | HC | 2,155 | *F* = 3.162 | **0.045** |
|  |  | Day | 1,155 | *F* = 3.327 | 0.07 |
|  |  | HC × Day | 2,155 | *F* = 1.652 | 0.195 |
|  |  | **Post hoc contrasts** |  |  |  |
|  |  | Low HC vs Ctrl (d2) | 151 | *t* = 2.170 | **0.032** |
|  |  | High HC vs Ctrl (d2) | 151 | *t* = −0.561 | 0.575 |
|  |  | Low HC vs Ctrl (d5) | 151 | *t* = 0.620 | 0.536 |
|  |  | High HC vs Ctrl (d5) | 151 | *t* = −0.150 | 0.881 |
|  |  | Low HC vs Ctrl (d8) | 151 | *t* = −0.160 | 0.873 |
|  |  | High HC vs Ctrl (d8) | 151 | *t* = −0.758 | 0.45 |

As behavioral changes were most pronounced on day 2, we summarized the phenotype at this time point using a radial plot (Figure 1D). Lower-HC exposure induced both anxiety-like behavior and hypolocomotion, whereas higher-HC exposure primarily impaired locomotion without a comparable increase in anxiety-like behavior. The anxiety-like, but not locomotor, effects of lower-HC exposure were replicated in an independent cohort (Supplementary Figure 1E–J). Together, these findings indicate that HC exposure produces its strongest behavioral effects after a single day of treatment and that stress intensity differentially shapes anxiety-like behavior and locomotion.

### Experiment 2.1: Lower-, but not higher-HC-induced anxiety is mediated by the serotonergic DRN

In Experiment 2, we investigated the role of the serotonergic dorsal raphe nucleus (DRN) in the anxiety-like phenotypes induced by different stress intensities. To this end, we repeated the day 1 HC treatment regimen from Experiment 1 (0, 0.5, and 1 μM), followed by chemogenetic ablation of the DRN overnight and behavioral testing on the subsequent day. To further characterize the behavioral effects of HC treatment, fish were assessed in both the standard SPM and a more aversive version of the test featuring increased illumination (Figure 2A).

**Figure 2.**
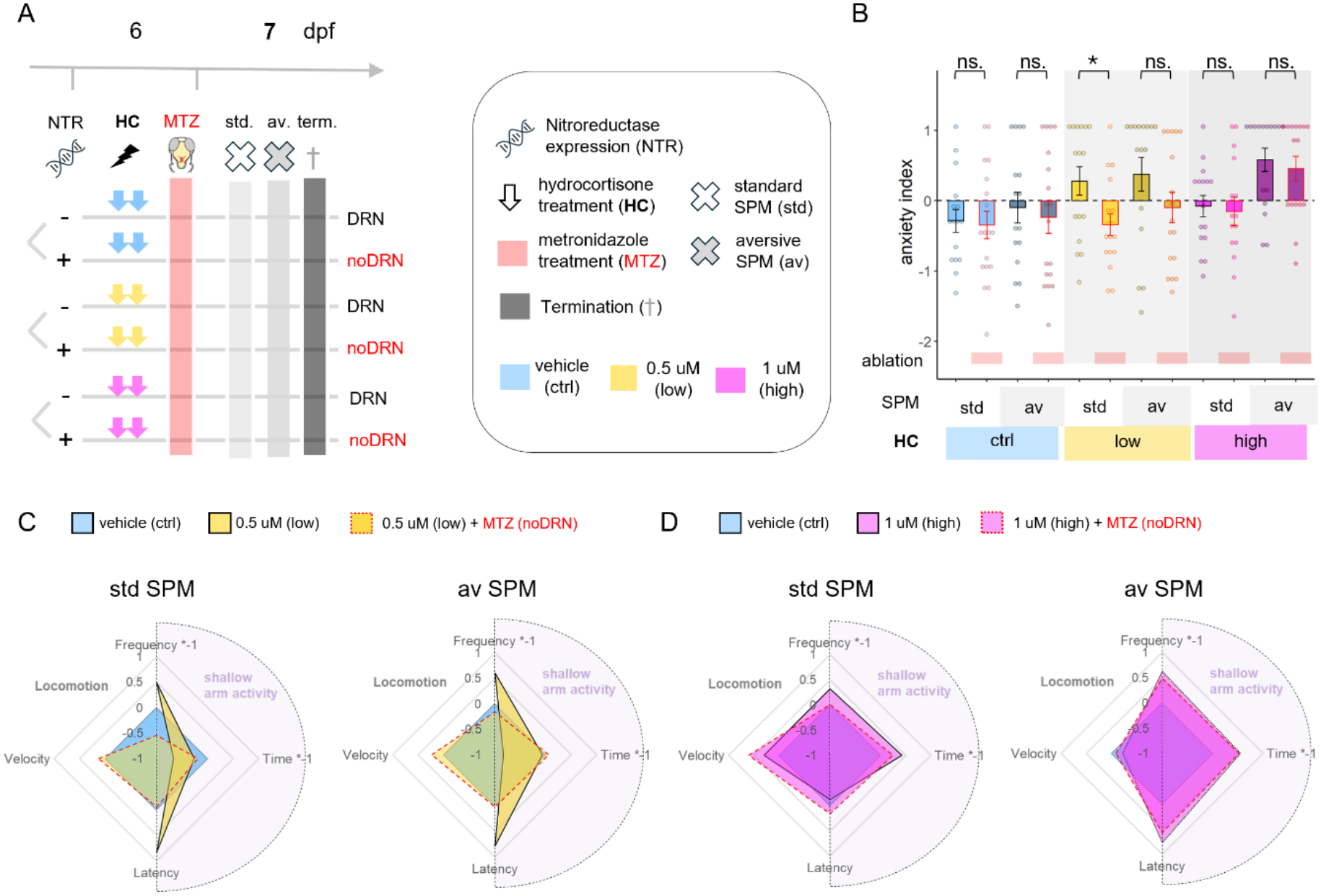
Stress-experience-dependent effects of DRN ablation on anxiety and locomotion. **A)** Treatment and sampling design of Experiment 2. **B)** Anxiety score ((shallow arm entries *-1 + shallow arm time *-1 + shallow arm latency)/3) measured in standard (std) and high aversivity (av) SPM following hydrocortisone (no, low and high stress) and MTZ (sham or DRN ablation) treatments. Red borders and rectangles indicate DRN ablated groups. Different tones of similar colors indicate type/aversivity of the SPM test. **C-D)** Radial plot of the behavioral phenotype. Mean velocity and variables of shallow arm activity (source variables of the anxiety score) in low stress **(C)** and high stress **(D)** HC groups measured in standard (std, on the left) and high aversivity (av, on the right) SPM tests. Note that shallow arm entries (frequency) and time is multiplied by −1 so that all shallow arm variables positively reflects anxiety. Variables are scaled and normed to the ctrl group. n=17(HC ctrl / DRN), 18 (HC ctrl / noDRN), 16 (HC low / DRN), 17 (HC low / noDRN), 16 (HC high / DRN), 16 (HC high / noDRN). Abbreviations: NTR (nitroreductase), HC (hydrocortisone), MTZ (metronidazole), std. (standard swimming plus-maze), av. (aversive swimming plus-maze), term (termination).

Increasing test illumination elevated anxiety-like behavior and reduced locomotion across all treatment groups, independent of DRN ablation, supporting that this test condition represents a different level of aversivity (Figure 2B, Supplementary Figure 2D, Table 2). Consistent with our previous findings, lower-HC exposure increased anxiety-like behavior in a standard SPM test, and furthermore, had a similar effect in the aversive SPM. In contrast, an anxiogenic effect of the higher-HC exposure only emerged under the more aversive testing conditions, indicating a context-dependent phenotype (Figure 2B, Supplementary Figure 2E).

**Table 2:** Statistical summary of Experiment 2.

| **Exp.** | **Dependent variable** | **Analysis / Effect** | **df** | **Statistic** | ***P*** |
|---|---|---|---|---|---|
| **2** | **Anxiety index** | **Three-way ANOVA** |  |  |  |
|  |  | HC | 2,185 | *F* = 5.553 | **0.005** |
|  |  | Ablation | 1,185 | *F* = 4.727 | **0.031** |
|  |  | SPM type | 1,185 | *F* = 7.719 | **0.006** |
|  |  | HC × Ablation | 2,185 | *F* = 1.783 | 0.171 |
|  |  | HC × SPM type | 2,185 | *F* = 1.950 | 0.145 |
|  |  | Ablation × SPM type | 1,185 | *F* = 0.001 | 0.979 |
|  |  | HC × Ablation × SPM type | 2,185 | *F* = 0.102 | 0.903 |
|  |  | **Post hoc contrasts** |  |  |  |
|  |  | noDRN vs DRN, Ctrl, low SPM | 185 | *t* = −0.207 | 0.836 |
|  |  | noDRN vs DRN, low HC, low SPM | 185 | *t* = −2.254 | **0.025** |
|  |  | noDRN vs DRN, high HC, low SPM | 185 | *t* = −0.262 | 0.794 |
|  |  | noDRN vs DRN, Ctrl, high SPM | 185 | *t* = −0.505 | 0.614 |
|  |  | noDRN vs DRN, low HC, high SPM | 185 | *t* = −1.715 | 0.088 |
|  |  | noDRN vs DRN, high HC, high SPM | 185 | *t* = −0.444 | 0.658 |
|  |  | Low HC vs Ctrl, noDRN, high SPM | 185 | *t* = 0.518 | 0.605 |
|  |  | High HC vs Ctrl, noDRN, high SPM | 185 | *t* = 2.556 | **0.011** |

Notably, these two anxiety phenotypes differed in their dependence on the DRN. Ablation of the DRN rescued the anxiety-like phenotype induced by lower HC in both testing conditions but had no effect on the anxiety induced by higher HC exposure (Figure 2B, Supplementary Figure 2F). Thus, lower-intensity stress produced a DRN-dependent, context-general anxiety state, whereas higher-intensity stress induced a DRN-independent anxiety state that became apparent only under highly aversive conditions (Figure 2C, 2D). DRN-dependence of lower-HC-induced anxiety was also confirmed in an independent cohort of animals (Supplementary Figure 2I-2M). Individual SPM measures are shown in Supplementary Figure 2 and Supplementary table 2.

Together, these findings indicate that the contribution of the DRN to anxiety depends on stress intensity and reveal at least two mechanistically distinct forms of stress-induced anxiety.

### Experiment 2.2: DRN-ablation-induced anxiolysis correlates with the activity of dopaminergic and oxytocinergic regions

To identify brain regions potentially involved in mediating the 5HT DRN-dependent anxiety in low HC treated groups, we compared phosphorylated ERK levels (pERK/tERK ratio) as a proxy for neuronal activity in the CNS following SPM testing in the same behaviorally characterized subjects as in Figure 2.

We first quantified neuronal activity across the 293 regions of the Z-Brain atlas^31^. Both HC treatments and DRN ablation produced widespread reductions in neuronal activity following SPM testing, as illustrated by representative activity maps (Figure 3A), maps of voxel-wise significant differences following DRN ablation (Figure 3B), and Linear Mixed Model (LMM) analysis of regional pERK activity (Figure 3C, Table 3). LMM analysis further revealed that DRN ablation significantly reduced activity in 56%, 14%, and 6% of CNS regions in the control, lower-HC, and higher-HC groups, respectively (Figure 3D). Most remaining regions were unaffected, whereas only 3% showed increased activity selectively in the lower-HC group.

**Figure 3.**
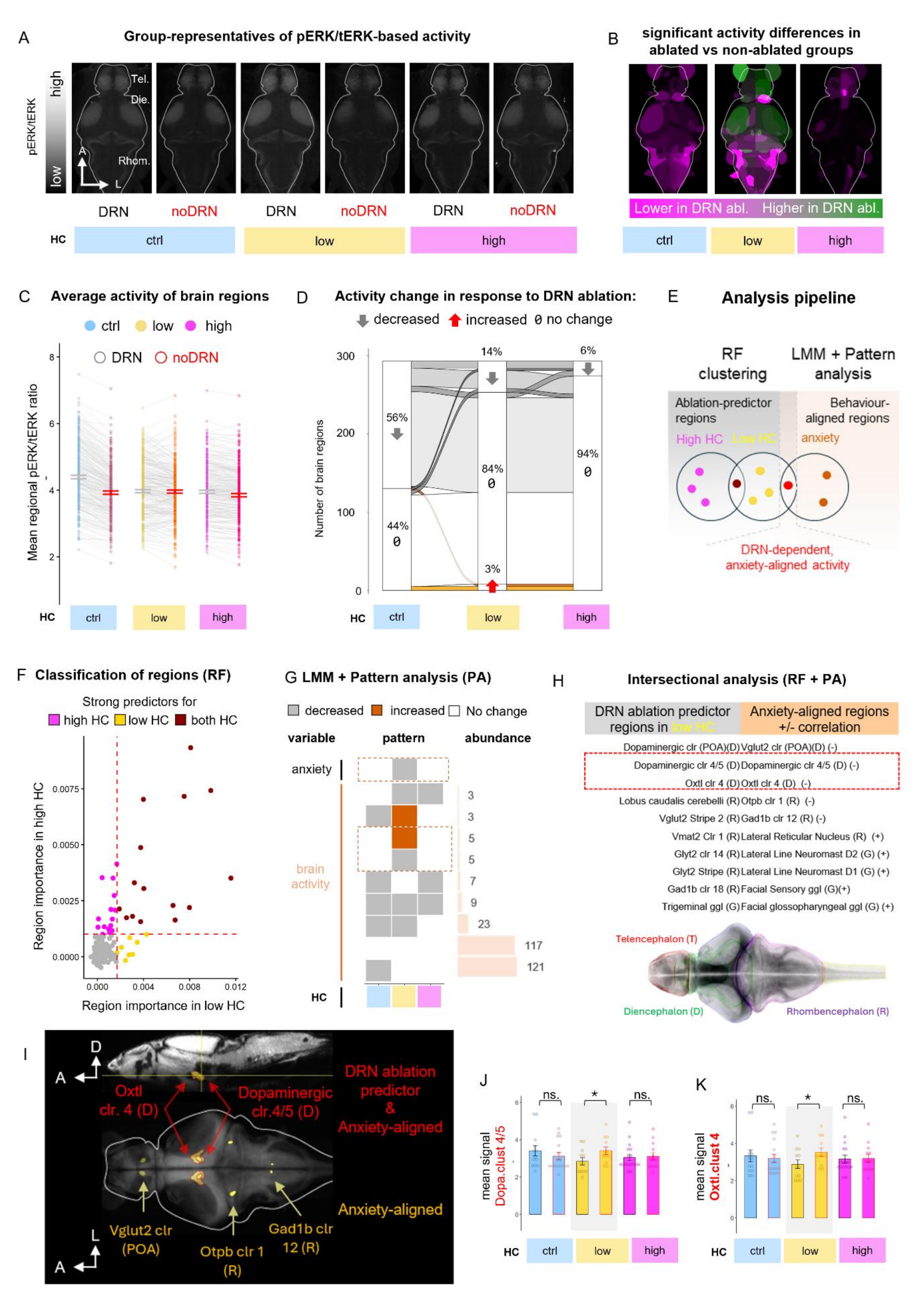
Stress-experience-dependent effects of DRN ablation on brain activity. **A)** Representative images of brain activity based on pERK/tERK ratio following hydrocortisone (ctrl, low and high stress) and MTZ (sham or DRN ablation) treatments. **B)** Z-brain regions decreased (magenta) or increased (green) their activity in response to DRN ablation following hydrocortisone (ctrl, low and high stress) treatment based on MAP-MAP analysis. **C)** Average activity (mean pERK/tERK signal) of assessed Z-brain brain regions. Each individual line represents a brain region. Red borders indicate DRN ablation (noDRN groups). **D)** Alluvial plot of activity change in response to ablation. Stacked columns represent the percentages of brain areas of different response types (decreased activity, increased activity, no change) to ablation based on the result of a linear mixed model and post-hoc contrasts in each stress treatment group. Channels connecting response types between stress treatment groups consists of brain regions that similarly changed their activity among groups (ablation*stress treatment). The thickness of channels represents the proportion of a pattern. **E)**. Design of sorting anxiety-aligned brain regions with the combined use of Random Forest (RF) clustering, linear mixed models (LMM) and consecutive pattern analysis. **F)** Permutation importance of brain regions in clustering subjects to DRN or to noDRN groups. Axis X and Y show the permutation-based importance in clustering in low and high HC treatment groups, respectively. Yellow vs magenta dots represent above-threshold predictors that selectively appear either in low or high HC groups, respectively. Dark red dots show shared predictors of ablation that appear in both HC groups. **G)** Pattern analysis based on the results of linear mixed models. Changes in anxiety or in brain activity in response to DRN ablation are indicated by grey (decrease), orange (increase) or blank (no change) squares in ctrl, low and high HC conditions. The abundance (number of brain regions) of each pattern is indicated on the right. Red dotted line circles the pattern of changes in anxiety and anxiety-correlated regions, showing either a negative or a positive relationship. **H)** Brain regions of low-HC selective, ablation-responsive cluster (left) and the anxiety-aligned cluster (right). Regions in the intersection of the two clusters are circled with red dashed line. **I)** Anxiolysis-aligned regions (orange/red) and those which are also strong predictors of DRN ablation (red) represented on a reference brain. **J-K)** Average activity (mean pERK/tERK signal) of the dopaminergic cluster 4/5 and the oxytocinergic cluster 4. Abbreviations: HC (hydrocortisone)

**Table 3:** Statistical summary of Experiment 2.2.

| Exp. | Dependent variable | Analysis / Effect | df | Statistic | *P* |
|---|---|---|---|---|---|
| **Exp. 2.2** | **Brain activity (Control HC)** | **Linear mixed model** |  |  |  |
|  |  | Ablation | 1,28 | *F* = 4.322 | **0.047** |
|  |  | Brain region | 292,8174 | *F* = 154.273 | **<0.001** |
|  |  | Ablation × Brain region | 292,8174 | *F* = 3.008 | **<0.001** |
| **Exp. 2.2** | **Brain activity (Low HC)** | **Linear mixed model** |  |  |  |
|  |  | Ablation | 1,22 | *F* = 0.403 | 0.532 |
|  |  | Brain region | 292,6413 | *F* = 87.833 | **<0.001** |
|  |  | Ablation × Brain region | 292,6413 | *F* = 3.598 | **<0.001** |
| **Exp. 2.2** | **Brain activity (High HC)** | **Linear mixed model** |  |  |  |
|  |  | Ablation | 1,28 | *F* = 0.561 | 0.46 |
|  |  | Brain region | 292,8153 | *F* = 136.551 | **<0.001** |
|  |  | Ablation × Brain region | 292,8153 | *F* = 1.299 | **<0.001** |

To identify regions that are potentially involved in the DRN-dependent anxiety induced by low HC treatment, we used an analysis pipeline combining a Random Forest analysis to determine regions predictive of DRN-ablation in the low HC treatment group, with an activity profile analysis to select regions that tracked the anxiety-like behavior profile across treatment groups (Figure 3E).

RF classification identified brain regions that predicted DRN ablation either across all HC treatment groups or selectively within a single HC condition (Figure 3F). Inflection-point analysis identified 30 lower-HC-selective and 26 higher-HC-selective predictor regions (Supp Fig 3 A-B).

Because RF identifies DRN-ablation dependent regions regardless of their behavioral relevance, we complemented this analysis with activity profile analysis to identify regional activity changes that specifically tracked the anxiolytic effect of DRN ablation across HC treatment groups. Based on region-specific change in activity in responses to DRN ablation across HC treatment groups, we identified nine distinct activity profiles (Figure 3G), two of which were positively, or negatively, associated with the change in anxiety-like behavior profiles across treatment groups. These two activity profiles were displayed by 10 anxiety-aligned CNS regions, which harbored selective changes in pERK levels selectively in low HC treatment (Fig 3 G,H). Among the 10 regions whose activity profiles mirrored the anxiolytic effect of DRN ablation, two were also among the strongest predictors of DRN ablation selectively in the lower-HC condition (Fig 3H): the dopaminergic cluster 4/5, which is located in the posterior tuberculum and hypothalamus, and the neighboring oxytocinergic Oxtl cluster 4, which is located in the hypothalamus (Fig 3 I). Comparison of mean pERK levels in these brain regions confirms the selective effect of DRN ablation on activity only in low HC group (Fig 3 J-K).

### Experiment 3: DRN-ablation-induced anxiolysis requires D1 receptor signaling and can be mimicked by dopamine reuptake inhibition

Given the anxiety-associated activity profile of dopaminergic cluster 4/5, we next investigated whether dopaminergic signaling mediates the anxiolytic effect of DRN ablation. Specifically, we asked (1) whether DRN-ablation-induced anxiolysis requires dopamine signaling and (2) which dopamine receptor subtype(s) mediate this effect. Fish were exposed to low-HC or vehicle treatment, underwent ablation or sham treatment overnight, and received systemic administration of either the D1 receptor antagonist SCH23390, or the D2 receptor antagonist haloperidol, the day after, prior to SPM testing (Figure 4A).

**Figure 4.**
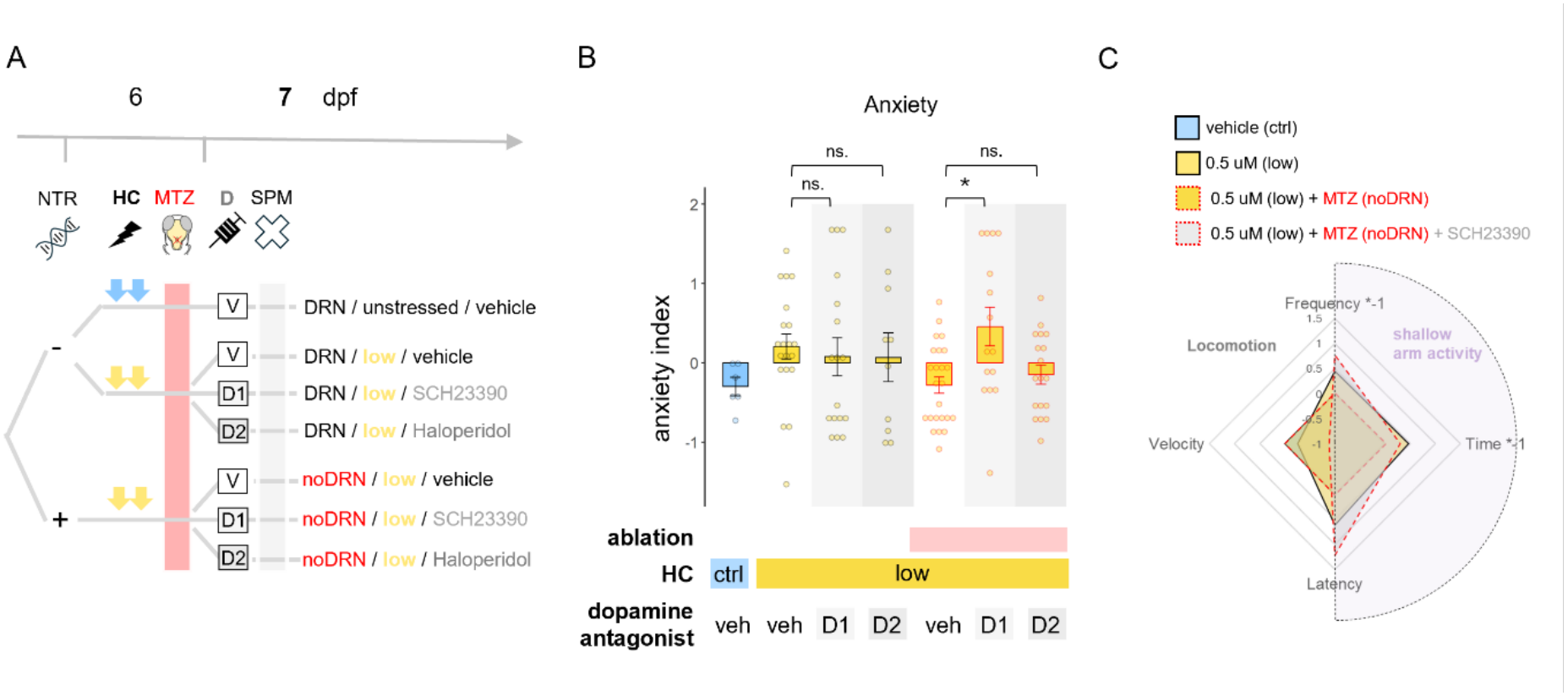
Dopaminergic mediators of serotonergic anxiety. **A)** Treatment and sampling design of Experiment 3. **B)** Anxiety score (SPM) following HC treatment and/or ablation and the acute application of vehicle, SCH23390 or haloperidol. **C)** Radial plot of the behavioral phenotype. Mean velocity and variables of shallow arm activity (source variables of the anxiety score) in low stress followed by sham/or DRN ablation and SCH23390 treatment. Note that shallow arm entries (frequency) and time is multiplied by −1 so that all shallow arm variables positively reflects anxiety. Variables are scaled and normed to the ctrl group. n=6(HC ctrl / DRN / D vehicle), 20 (HC low / DRN / D vehicle), 17 (HC low / DRN / D D1), 10 (HC low / DRN / D D2), 24 (HC low / noDRN / D vehicle), 15 (HC low / noDRN / D D1), 18 (HC low / noDRN / D D2) Abbreviations: NTR (nitroreductase), HC (hydrocortisone), MTZ (metronidazole), SPM (swimming plus-maze), D (dopaminergic treatment), V (vehicle), D1 (SCH23390), D2 (haloperidol).

As observed previously, we confirmed that DRN ablation reduced the anxiety-like behavior induced by low-HC treatment. This anxiolytic effect of DRN ablation was abolished by D1, but not D2, receptor blockade (Figure 4B, Table 4). Interestingly, neither receptor antagonist influenced anxiety in non-ablated, DRN intact conditions. D1 receptor blockade reduced swimming speed in both ablated and non-ablated fish, with a smaller effect in the ablated group (Supplementary Figure 4D, Supplementary table 4). Together, these findings demonstrate that D1 receptor signaling is required, potentially as a critical downstream mediator, for the anxiolytic effect of DRN ablation.

**Table 4:** Statistical summary of Experiments 3.

| **Exp.** | **Dependent variable** | **Analysis / Effect** | **df** | **Statistic** | ***P*** |
|---|---|---|---|---|---|
| **Exp. 3** | **Anxiety index** | **Three-way ANOVA** |  |  |  |
|  |  | Ablation | 1,103 | *F* = 1.437 | 0.233 |
|  |  | Dopamine antagonist | 2,103 | *F* = 1.809 | 0.169 |
|  |  | Ablation × Dopamine antagonist | 2,103 | *F* = 3.178 | **0.046** |
|  |  | **Post hoc contrasts** |  |  |  |
|  |  | D1 vs Vehicle, DRN | 103 | *t* = −0.529 | 0.598 |
|  |  | D2 vs Vehicle, DRN | 103 | *t* = −0.481 | 0.631 |
|  |  | D1 vs Vehicle, noDRN | 103 | *t* = 3.017 | **0.003** |
|  |  | D2 vs Vehicle, noDRN | 103 | *t* = 0.579 | 0.564 |

## DISCUSSION

Anxiety disorders show marked variability in both pathophysiology and treatment response, yet the mechanisms underlying this heterogeneity remain poorly understood. Here, we demonstrate that stress intensity determines whether anxiety recruits serotonergic mechanisms. Specifically, lower HC exposure produced a DRN-dependent anxiety state, whereas higher HC exposure generated a mechanistically distinct DRN-independent anxiety state. We further identify D1 receptor-dependent dopaminergic signaling as a downstream mediator of the serotonergic anxiety state.

### Stress intensity determines the form of anxiety

Previous studies have shown that the behavioral consequences of stress depend on the controllability, modality^17,24^, duration^24^, and temporal pattern (i.e. interrupted vs continuous presentation)^33^ of the stressor. Our findings extend this concept by identifying stressor intensity as an additional determinant of anxiety expression. Notably, increasing the concentration of hydrocortisone exposure did not simply enhance anxiety but shifted its expression from a context-general to a context-dependent phenotype, suggesting that stronger stress alters the threshold at which anxiety is behaviorally expressed.

Our observation that anxiety was induced by an intermediate, but not higher, glucocorticoid concentration in the standard SPM is consistent with extensive evidence for non-linear effects of stress and glucocorticoids on behavior. Stress magnitude and glucocorticoid levels frequently show inverted-U-shaped relationships with performance across behavioral domains. Ryu and colleagues systematically manipulated stress intensity and found that moderate, but not higher, stress enhanced innate responses including optomotor behavior, rheotaxis, and heat avoidance. These non-linear changes in behavioral responsiveness were inversely mirrored by subjects’ cortisol levels^34^. One explanation for such non-linearity is that different stress intensities recruit distinct glucocorticoid-dependent processes. Different stressors can produce qualitatively distinct defensive phenotypes in zebrafish, ranging from reduced exploration to freezing and erratic swimming^35^. In line with this, lower glucocorticoid levels preferentially engage mineralocorticoid receptor-dependent processes associated with anxiety, whereas higher levels increasingly recruit glucocorticoid receptor-dependent mechanisms involved in contextual processing and memory^36^. Differential recruitment of these pathways may therefore contribute to the distinct anxiety phenotypes observed following lower versus higher HC exposure.

Alternatively, different stress intensities may recalibrate the allostatic set-point of stress reactivity, shifting the threshold for behavioral expression such that anxiety emerges in one environmental context but not another, rather than simply producing quantitatively different levels of anxiety. Cheng and colleagues demonstrated that combinations of osmotic stress intensity and test-aversiveness recruit distinct anxiety-related behavioral domains in larval zebrafish, with stronger stress reducing the correlation between these domains^37^. In our previous work, we similarly found that social isolation altered behavioral responsiveness in a testing-environment-dependent manner, producing different effects under baseline and novel conditions. Prior stress has also been shown to reprogram subsequent stress responsiveness at the level of cortisol dynamics^38^. Together, these findings raise the possibility that lower HC more directly promotes anxiety expression, whereas higher HC recalibrates the conditions under which anxiety is subsequently expressed, resulting in the context-dependent phenotype observed here.

### Stress intensity determines the involvement of the serotonergic DRN in anxiety

Here, we show that of the two stress-intensity-dependent anxiety phenotypes, only lower-HC-induced anxiety involved the serotonergic DRN. Although previous studies have not directly examined whether stress intensity determines serotonergic involvement in anxiety, other properties of prior stress appear to do so. We previously showed that both social isolation and chronic unpredictable stress (CUS) perturb the serotonergic system while increasing anxiety. However, whereas isolation-induced serotonergic changes and anxiety were rescued by the serotonergic anxiolytic buspirone, CUS-induced anxiety persisted following ablation of serotonergic DRN neurons^11,21^. Differences in serotonergic involvement between social and non-social stressors have also been reported across species^17,24^, while stressor duration has been identified as another important determinant^10^. Our findings extend this framework by identifying stressor intensity as an additional factor determining serotonergic involvement in anxiety regulation.

A potentially overlapping explanation is that serotonergic signaling preferentially contributes to context-general rather than context-specific defensive behavior. Hui-quan Li et al. demonstrated that generalized – but not context-specific – fear induced by either intense electric shock or glucocorticoid treatment was associated with a neurotransmitter switch from glutamate to GABA in dorsal raphe serotonergic neurons and could be prevented by early SSRI administration^39^. Although generalized fear is distinct from anxiety, the two represent related defensive states and may share mechanisms regulating the context dependence of their expression^40^. Consistent with a broader role in generalized anxiety, serotonergic agents originally developed for major depression, including SSRIs and the 5-HT1A receptor partial agonist buspirone, have become established treatments for generalized anxiety disorder.

Together with our previous work^11,21^, our findings suggest that serotonergic involvement is not an intrinsic feature of anxiety itself but depends on the stress history from which anxiety emerges. In our model, the context-general anxiety state induced by lower HC required the DRN for its expression, whereas the context-dependent anxiety induced by higher HC did not. Thus, stressor intensity and the resulting context dependence of anxiety may represent interconnected determinants of serotonergic involvement in anxiety regulation.

### A downstream dopaminergic component of DRN-dependent anxiety

Our whole-brain activity analysis identified dopaminergic cluster 4/5 of the Z-Brain atlas as a candidate region associated with DRN-dependent anxiolysis. Although this analysis does not establish a causal role in anxiolysis for this population, it suggests that dopaminergic circuits may contribute to the downstream effects of the DRN. Consistent with this hypothesis, D1 receptor blockade abolished the anxiolytic effect of DRN ablation in our model.

Previous studies have similarly implicated dopaminergic signaling in anxiety regulation. Patients with generalized^41^ and social anxiety^42^ disorders exhibit reduced dopamine transporter availability, suggesting diminished dopaminergic tone. In rodents, threatening stimuli suppress the activity of midbrain dopaminergic neurons, whereas similar location neurons become more active during exploration of aversive environments^43,44^.

D1 receptor signaling has likewise been repeatedly linked to anxiety, although its effects appear to depend strongly on the anatomical circuit and behavioral context^45–48^. While systemic43, cortical^49^, or amygdalar^45^ D1 receptor antagonism has produced anxiolytic effects in some studies, antagonism within the midbrain^50^ or striatum^47^ is predominantly anxiogenic. Similarly, D1 receptor blockade abolishes the anxiolytic effects of both adenylyl cyclase 5 deficiency^48^ and neurotensin administration^50^, further supporting a context-dependent role of D1 signaling in anxiety regulation.

### Conclusion

Together, our findings identify stress intensity as a previously unrecognized determinant of serotonergic recruitment during anxiety and support a downstream contribution of dopaminergic signaling to DRN-dependent anxiety. By demonstrating that distinct stress histories recruit different neurochemical mechanisms, our study provides a framework for understanding the biological heterogeneity of anxiety and the variable efficacy of serotonergic treatments.

## MATERIALS AND METHODS

### Animals

Experiments were performed using larval transgenic nacre mifta -/- zebrafish^51^ (*Danio rerio*) carrying the Tg(tph2:Gal4; UAS:nitroreductase-mCherry) transgene and aged 6–16 days post fertilization (dpf). This transgenic line expresses the bacterial nitroreductase (NTR) enzyme together with mCherry selectively in serotonergic neurons of the dorsal raphe nucleus (DRN), enabling their chemogenetic ablation following metronidazole treatment^52^.

Fish were maintained at 28°C under a 10 h light/14 h dark photoperiod and fed commercial flake food (TetraMin) twice daily. Larvae were group-housed (40–60 fish per tank) in mesh-bottom nursery tanks within a recirculating aquatic system (Techniplast). All experimental procedures were approved by the Danish Animal Experiments Inspectorate (permit no. 2023-15-0201-01493).

### Hydrocortisone treatment

Hydrocortisone (HC) treatment at different concentrations was used to model different stress intensities. Fish were exposed to either 0.5 μM (low-HC) or 1 μM (high-HC) hydrocortisone, representing lower- and higher-intensity HC treatment, respectively. Concentrations were selected based on previous studies^53^ and experiments performed in our laboratory^54^.

Hydrocortisone (H4881-1G, Sigma-Aldrich) was prepared as 1 mM stock aliquots in fish water, stored at −18°C, and freshly diluted in fish water to the required working concentration immediately before each treatment. Fish received two 20-minute HC exposures daily, each followed by a 20-minute washout in fish water. In Experiment 1, treatments were administered daily from 6 to 13 dpf. In all subsequent experiments, fish received HC treatment only at 6 dpf.

### Chemogenetic DRN ablation

Serotonergic neurons of the dorsal raphe nucleus (DRN) were selectively ablated using the nitroreductase/metronidazole (NTR/MTZ) system. Nitroreductase-expressing Tg(*tph2:Gal4; UAS:nitroreductase-mCherry*) larvae were exposed to 5 mM metronidazole (MTZ; Merck) dissolved in fish water, whereas their mCherry-negative siblings received the same MTZ treatment as sham controls.

Fish were exposed to MTZ overnight before behavioral testing. During MTZ exposure, larvae were maintained in 200 mL aluminum foil-covered Petri dishes to protect the light-sensitive compound from degradation. The ablation protocol was optimized and validated in our previous study^11^.

### Drugs

Haloperidol and SCH23390 (Sigma-Aldrich) were dissolved in 0.01% dimethyl sulfoxide (DMSO) and prepared as 0.1 mM stock solutions. Stock solutions were stored as aliquots at −20°C and freshly diluted to their working concentrations immediately before each experiment. The final concentrations used were 1 µM haloperidol and 10 µM SCH23390.

### Swimming plus-maze test

The swimming plus-maze (SPM) is a high-throughput behavioral assay that measures anxiety-like surface avoidance in larval zebrafish developed by ZKV^28^. The apparatus consisted of a plus-shaped platform with two shallow (length x width x depth: 10 × 8 × 2.5 mm) and two deep (length x width x depth : 10 × 8 × 5 mm) arms connected by a central zone. To manipulate the aversiveness of the testing environment, platforms were 3D printed from either white resin (standard SPM) or translucent resin (aversive SPM).

Behavioral testing was conducted in a dark room under uniform bottom illumination provided through a diffuser by a white and an infrared LED light sources and. All behavioral tests were conducted at 28 °C in a temperature-controlled room, after 1 pm. Individual fish were transferred from their home tank to the center of the maze, and swimming behavior was recorded for 6 min at 5 frames. s⁻¹ using a Basler ace2 camera fitted with a 35 mm lens and a long-pass infrared filter. The setup consisted of 12 adjacent SPM platforms so that 12 fish were recorded simultaneously. Anxiety-like behavior was quantified as the time spent in, number and first latency of entries to the shallow arms, while mean swimming velocity was used as a measure of locomotor activity.

### Immunohistochemistry

Because ERK is rapidly phosphorylated following neuronal activation, brain activity during the SPM testing was assessed by immunohistochemical detection of phosphorylated and total extracellular signal-regulated kinase (pERK and tERK, respectively) in all experiments. The staining protocol was adapted from Randlett *et al.*^31^.

Immediately after behavioral testing, larvae were euthanized and fixed overnight at 4°C in ice-cold 10% PFAT (10% paraformaldehyde containing 0.25% Triton X-100). Samples were subsequently stored in PBST (PBS containing 0.25% Triton X-100) at 4°C for at least one month to ensure adequate tissue permeabilization.

For immunostaining, samples were permeabilized in 0.05% Trypsin-EDTA (25200-056, Thermo Fisher Scientific) for 45 min on ice with gentle agitation, washed three times for 5 min in PBST, and incubated for 1 h at room temperature in blocking solution containing 2% normal goat serum (NGS), 1% bovine serum albumin (BSA; A9647, Sigma-Aldrich), and 1% dimethyl sulfoxide (DMSO; D8418, Sigma-Aldrich) prepared in PBST.

Primary antibodies were diluted in blocking solution and applied for 72 h at 4°C (mouse anti-ERK, L34F12, Cell Signaling Technology, 1:300; rabbit anti-pERK, D13.14.4E, Cell Signaling Technology, 1:600). Following five 5-min washes in PBST, samples were incubated overnight at 4°C with Alexa Fluor 633 goat anti-mouse IgG and Alexa Fluor 488 goat anti-rabbit IgG secondary antibodies (Thermo Fisher Scientific; both 1:600). After three additional 5-min washes in PBST, samples were stored at 4°C until confocal imaging.

### Confocal microscopy

Before imaging, samples were washed in artificial fish water (AFW) and mounted ventral side down in 2% low-melting-point agarose within silicone imaging chambers on microscope slides. Whole-brain image stacks were acquired using a Zeiss LSM700 laser-scanning confocal microscope (Carl Zeiss AG) equipped with a W N-Achroplan 10×/0.3 W M27 objective. Images were acquired using 488 nm and 639 nm diode lasers at 16-bit depth with an xy resolution of 0.63 µm pixel⁻¹ and a z-step of 7.18 µm. The laser intensity, gain and zoom factors were kept constant, and image stacks were acquired in LineSequential mode.

### Experimental design

#### Experiment 1

To establish a hydrocortisone (HC)-based model of graded stress exposure, larvae received vehicle, 0.5 µM HC (low-HC), or 1 µM HC (high-HC) twice daily from 6 to 13 dpf. Behavioral testing was performed before treatment (6 dpf) and subsequently at 8, 10, 12, and 14 dpf using separate sets of fish at each time point. An additional independent cohort was included to replicate the behavioral effects observed following a single day of low-HC exposure.

#### Experiment 2

To investigate the contribution of the dorsal raphe nucleus (DRN) to HC-induced anxiety, fish were assigned to six experimental groups comprising DRN-intact and DRN-ablated larvae exposed to vehicle, low-HC (0.5 µM), or high-HC (1 µM). Following genotype-based sorting, larvae received two 20-min HC treatments at 6 dpf, followed by chemogenetic DRN ablation overnight. Behavioral testing was performed the following day using the swimming plus-maze (standard followed by aversive configuration). Whole-body samples were collected immediately after behavioral testing for pERK/tERK immunohistochemistry. An additional independent cohort was included to replicate the behavioral effects of DRN ablation following low-HC treatment.

#### Experiment 3

To determine whether dopamine receptor signaling mediates the anxiolytic effects of DRN ablation, larvae were assigned to seven experimental groups: DRN-intact control + vehicle, DRN-intact low-HC + vehicle, haloperidol, or SCH23390, and DRN-ablated low-HC + vehicle, haloperidol, or SCH23390. Larvae received two 20-min low-HC treatments at 6 dpf, followed by overnight DRN ablation. Behavioral testing was conducted in the standard swimming plus-maze the following day. Pharmacological treatments (vehicle, haloperidol, or SCH23390) were administered by bath application for 30 min before behavioral testing, followed by a 1-minute wash-out in 6 well-plates.

### Data analysis

#### Behavioral data

Sample sizes were estimated based on an expected effect size of 0.4 for anxiety-like behavior, derived from previous chronic unpredictable stress experiments, assuming a one-way ANOVA with three experimental groups.

All statistical analyses were performed in R (R Foundation for Statistical Computing, Vienna, Austria^55^). To obtain a composite measure of anxiety-like behavior in the swimming plus-maze (SPM), an anxiety index was calculated by combining three behavioral variables^11,54^: latency to enter the shallow arms, frequency of shallow arm entries, and time spent in the shallow arms. Individual variables were first scaled, after which the anxiety score was calculated as:

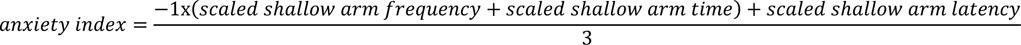

The anxiety index was used as the primary dependent variable in all analyses. The effects of hydrocortisone treatment were assessed using two-way ANOVA, whereas the combined effects of previous stress exposure, current stress conditions, and DRN ablation were analyzed using three-way ANOVA, including interaction terms. Pairwise comparisons were performed using estimated marginal means (emeans). In cases where the magnitude of group variability changed along with group differences, two-group comparisons were performed using the Brunner–Munzel test.

Data are presented as mean ± standard error of the mean (SEM). Statistical significance was defined as *P* < 0.05.

#### Whole-brain activity mapping

Whole-brain neuronal activity mapping (MAP-map) was performed using a modified version of the pipeline developed by Randlett *et al*.^31^. The analysis comprised image registration, voxel-wise comparison of pERK/tERK activity, and anatomical quantification.

#### Image registration and preprocessing

Individual brain scans were registered using the Computational Morphometry Toolkit (CMTK)^56^ to a 7 dpf reference brain. Initial registrations were performed by registering the tERK channel of each fish against the reference brain of the zbrain atlas using the CMTK command-line interface (parameters: ߢ-T 16 -X 52 -C 8 -G 80 -R 3 -A ‘--accuracy 0.4’ -W ‘-- accuracy 1.6’’). The four highest-quality registrations were used to generate a study-specific average tERK template, to which all remaining brains were subsequently registered. Registered image stacks were preprocessed in FIJI/ImageJ^57,58^ using the PrepareStacksForMAPMapping.ijm macro^31^, including resampling (300 × 679 × 80) and two-dimensional (xy) Gaussian filtering.

#### MAP-map generation

Voxel-wise comparisons of pERK/tERK activity were performed in MATLAB using the MakeTheMAPMap.m function from the MAP-mapping pipeline^31^. Group differences were quantified using Mann–Whitney U-based Z-scores. Statistical significance was determined using false discovery rate (FDR) correction, with voxels exceeding the FDR threshold (0.005% false-positive rate in control voxels) considered significantly different. Significant voxels were assigned to intensity values corresponding to the median difference between groups (scaled from 0 to 65,535) and color-coded according to the direction of change (green, increased activity; magenta, decreased activity).

#### Regional quantification

To quantify activity within each of the 293 regions of the Zbrain atlas, pERK and tERK average intensity were extracted from the registered brain scans using a custom-written Python script. Mean pERK/tERK ratii were calculated for each Z-Brain anatomical region across all animals.

#### Random Forest and pattern analysis

To identify brain regions responsive to dorsal raphe nucleus (DRN) ablation and determine which of these were specifically associated with anxiolysis, we combined Random Forest (RF) classification with linear mixed models and pattern analysis.

Random Forest is a permutation-based machine learning algorithm that identifies combinations of variables that best discriminate between predefined groups^59^. Separate RF models were constructed for the control vs low- and control vs high-HC treatment groups, each running 4000 permutations, to identify anatomical regions whose activity best distinguished DRN-ablated from DRN-intact larvae in each condition. Strongest predictors were chosen based on an inflection point analysis (inflection package) of permutation importance of variables (brain regions) (Supplementary Figure 3B). Regions identified by RF were subsequently classified as low-HC-specific, high-HC-specific, or shared between treatment groups.

To determine whether DRN-responsive regions tracked behavioral rescue, regional pERK/tERK activity was analyzed using linear mixed models with DRN ablation as the explanatory variable within each HC treatment. Based on the direction and significance of the ablation effect based on post hoc contrasts (emeans package), brain regions were grouped according to their activity patterns across treatment conditions. Regions showing selective activity changes associated with anxiolysis in the low-HC condition were distinguished from those exhibiting shared or HC-independent response patterns.

Finally, candidate substrates of DRN-dependent anxiolysis were identified as the intersection between regions strongly discriminating DRN ablation in the RF analysis and regions whose activity patterns tracked anxiolytic behavioral responses in the pattern analysis.

## AUTHORS CONTRIBUTIONS

Z.K.V & F.K. conceived the study and supervised; Z.K.V conducted the experiments, with help from J.V.M., L.J.F. & A.U.; Z.K.V analyzed the data, with some help from J.V.M; Z.K.V prepared the figures and wrote the original draft; F.K. and L.J.F. reviewed and edited the manuscript.

## ACKNOWLEDGMENTS

We thank Dr. Harold Burgess for generously sharing the transgenic Tg(Tph2:Gal4:UAS-Ntr-mCherry) line. We thank the expert staff from the Core Facility for Integrated Bioimaging (CFIM/CFIB) at UCPH for training and guidance with microscopy image acquisition. This work was funded by the Lundbeck Foundation (Ascending Investigator grant to F.K.).

## SUPPLEMETARY FIGURES

**Supplementary Figure 1.**
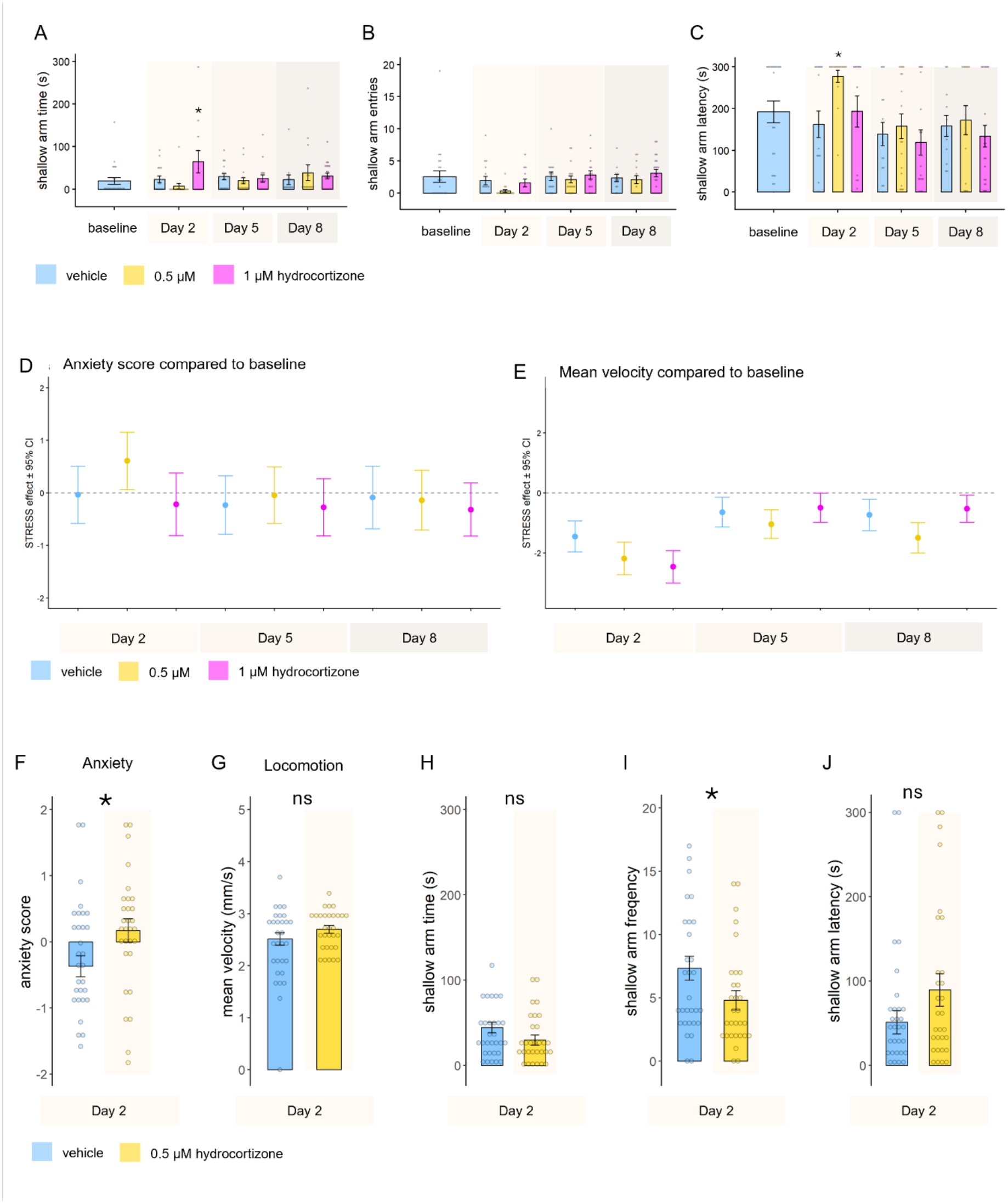
Concentration-and duration-dependent effects of hydrocortisone stress on anxiety and locomotion. **A-C)** Time spent in **(A)**, frequency **(B)** and latency **(C)** of entries to the shallow arms of the SPM before and after treatments of hydrocortisone. **D-E)** Confidence intervals of the hydrocortisone effect on anxiety **(D)** and mean velocity **(E)** compared to baseline sampling. **F-J)** Results of an independent confirmation experiment of HC-induced (0 vs 0.5 μM) anxiety on day 2 in the SPM. n=30 fish/ group.

**Supplementary Figure 2.**
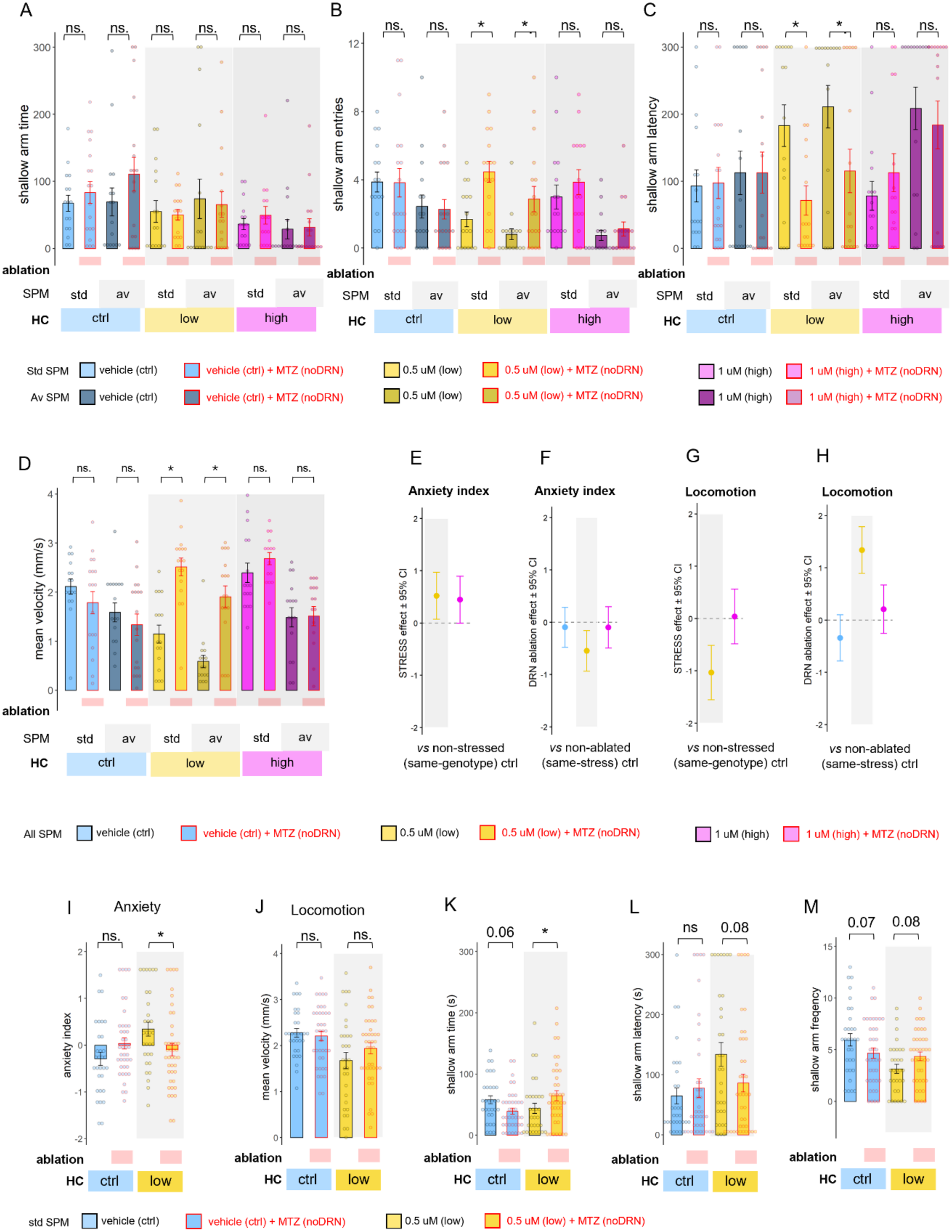
Stress-experience-dependent effects of DRN ablation on anxiety and locomotion. **A-D)** Time spent in **(A)**, frequency **(B)** and latency **(C)** of entries to the shallow arms, and mean swimming speed **(D)** measured in standard (std) and high aversivity (av) SPM following hydrocortisone (no, low and high stress) and MTZ (sham or DRN ablation) treatments. Red borders and rectangles indicate DRN ablated groups. Different tones of similar colors indicate type/aversivity of the SPM test. D-F) Anxiety index shown regardless of SPM type measured in all SPM following hydrocortisone (no, low and high stress) and MTZ (sham or DRN ablation) treatments. **E-F)** 95% confidence intervals of difference in anxiety index in stress vs corresponding (same genotype) unstressed groups **(E)** and ablated vs corresponding (same stress) non-ablated groups **(F)** regardless of SPM types. Group differences are considered significant if the error bars of the confidence interval do not cross the dotted line **G-H)** 95% confidence intervals of difference in mean swimming speed in stress vs corresponding (same genotype) unstressed groups **(G)** and ablated vs corresponding (same stress) non-ablated groups **(H)** regardless of SPM types. Group differences are considered significant if the error bars of the confidence interval do not cross the dotted line **D-H)** Results of an independent confirmation experiment of HC-induced (0 vs 0.5 μM) anxiety and DRN ablation-induced (DRN vs noDRN) anxiolysis on day 2 in the standard SPM. Fish numbers: n=31(HC ctrl/DRN), 39(HC ctrl/noDRN), 32(HC low/DRN), 41(HC low/noDRN).

**Supplementary Figure 3.**
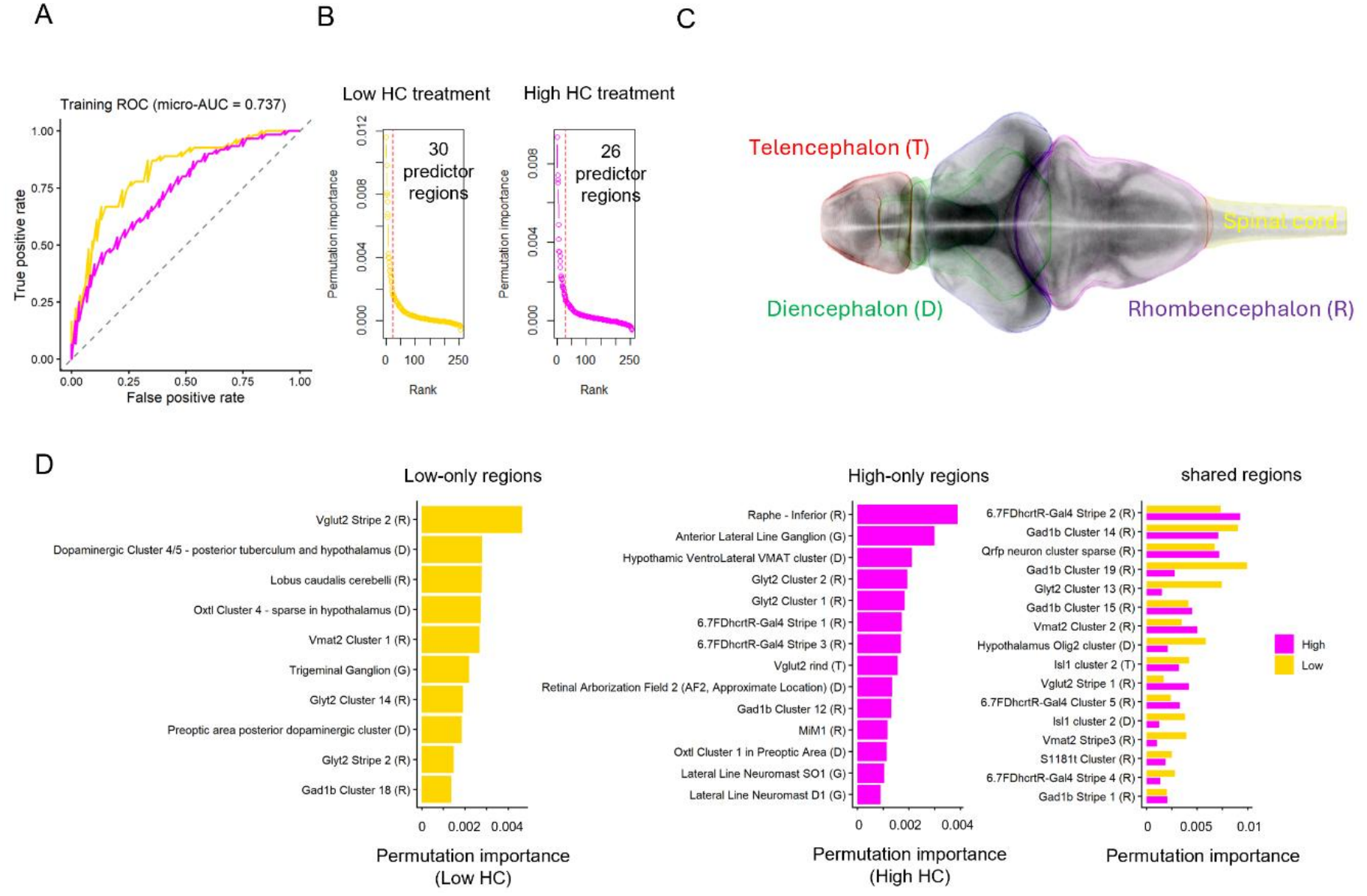
Stress-experience-dependent effects of DRN ablation on brain activity. **A)** ROC curve of clustering accuracy of Random Forest clustering analysis in low (gold) and high (magenta) HC treatment groups. **B)** Permutation importance of all assessed brain regions in classifying subjects to DRN and noDRN groups in low and high HC treatments. Red dashed lines indicate a threshold for significant predictors (left of the line) based on inflection point analysis. **C)** Delineation of main central nervous system subdivisions in the atlas. **D)** Most important predictor regions for DRN-ablation in low (left), high (middle) and both (right) HC treatments.

**Supplementary Figure 4.**
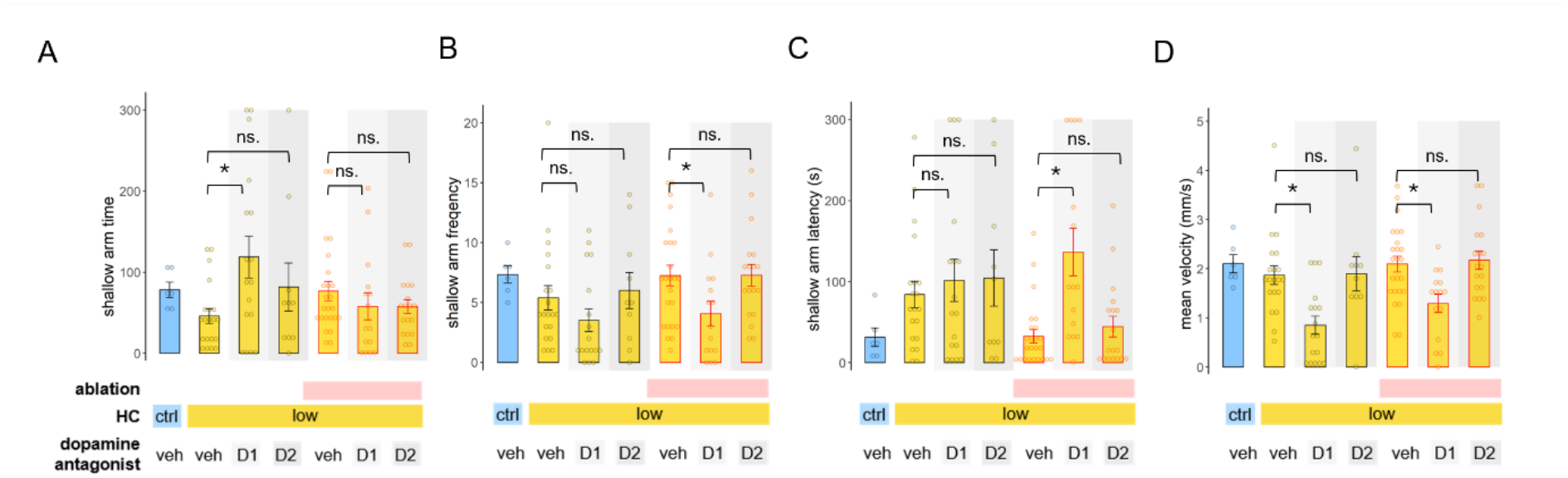
Dopaminergic mediators of serotonergic anxiety. **A–D)** Time spent in the shallow arm (A), number (B) and latency (C) of entries into the shallow arm, and mean swimming speed (D) in the SPM following HC treatment and/or DRN ablation and acute treatment with vehicle, SCH23390, or haloperidol. Abbreviations: NTR (nitroreductase), HC (hydrocortisone), MTZ (metronidazole), D (dopaminergic treatment), V (vehicle), D1 (SCH23390), D2 (haloperidol).

## SUPPLEMETARY TABLES

**Supplementary Table 1.**

| **Exp.** | **Dependent variable** | **Analysis / Effect** | **df** | **Statistic** | ***P*** |
|---|---|---|---|---|---|
| **1** | **Mean velocity** | **Two-way ANOVA** |  |  |  |
|  |  | HC | 2,144 | *F* = 12.616 | **<0.001** |
|  |  | Day | 1,144 | *F* = 8.921 | **0.003** |
|  |  | HC × Day | 2,144 | *F* = 12.571 | **<0.001** |
|  |  | **Post hoc contrasts** |  |  |  |
|  |  | Low HC vs Ctrl (d2) | 140 | *t* = −2.448 | **0.016** |
|  |  | High HC vs Ctrl (d2) | 140 | *t* = −3.367 | **<0.001** |
|  |  | Low HC vs Ctrl (d5) | 140 | *t* = −1.549 | 0.124 |
|  |  | High HC vs Ctrl (d5) | 140 | *t* = 0.579 | 0.563 |
|  |  | Low HC vs Ctrl (d8) | 140 | *t* = −2.657 | **0.009** |
|  |  | High HC vs Ctrl (d8) | 140 | *t* = 0.792 | 0.43 |
| **1** | **Shallow arm entries** | **Two-way ANOVA** |  |  |  |
|  |  | HC | 2,155 | *F* = 2.476 | 0.087 |
|  |  | Day | 1,155 | *F* = 3.277 | 0.072 |
|  |  | HC × Day | 2,155 | *F* = 1.228 | 0.296 |
|  |  | **Post hoc contrasts** |  |  |  |
|  |  | Low HC vs Ctrl (d2) | 151 | *t* = −1.725 | 0.087 |
|  |  | High HC vs Ctrl (d2) | 151 | *t* = −0.348 | 0.728 |
|  |  | Low HC vs Ctrl (d5) | 151 | *t* = −0.511 | 0.61 |
|  |  | High HC vs Ctrl (d5) | 151 | *t* = 0.222 | 0.825 |
| **Exp.** | **Dependent variable** | **Analysis / Effect** | **df** | **Statistic** | ***P*** |
|  |  | Low HC vs Ctrl (d8) | 151 | *t* = −0.250 | 0.803 |
|  |  | High HC vs Ctrl (d8) | 151 | *t* = 0.790 | 0.431 |
| **1** | **Shallow arm time** | **Two-way ANOVA** |  |  |  |
|  |  | HC | 2,155 | *F* = 2.056 | 0.131 |
|  |  | Day | 1,155 | *F* = 0.063 | 0.801 |
|  |  | HC × Day | 2,155 | *F* = 3.759 | **0.025** |
|  |  | **Post hoc contrasts** |  |  |  |
|  |  | Low HC vs Ctrl (d2) | 151 | *t* = −1.073 | 0.285 |
|  |  | High HC vs Ctrl (d2) | 151 | *t* = 2.485 | **0.014** |
|  |  | Low HC vs Ctrl (d5) | 151 | *t* = −0.639 | 0.524 |
|  |  | High HC vs Ctrl (d5) | 151 | *t* = −0.316 | 0.752 |
|  |  | Low HC vs Ctrl (d8) | 151 | *t* = 0.961 | 0.338 |
|  |  | High HC vs Ctrl (d8) | 151 | *t* = 0.574 | 0.567 |
| **1** | **Shallow arm latency** | **Two-way ANOVA** |  |  |  |
|  |  | HC | 2,155 | *F* = 3.172 | **0.045** |
|  |  | Day | 1,155 | *F* = 6.879 | **0.01** |
|  |  | HC × Day | 2,155 | *F* = 1.042 | 0.355 |
|  |  | **Post hoc contrasts** |  |  |  |
|  |  | Low HC vs Ctrl (d2) | 151 | *t* = 2.825 | **0.005** |
|  |  | High HC vs Ctrl (d2) | 151 | *t* = 0.705 | 0.482 |
|  |  | Low HC vs Ctrl (d5) | 151 | *t* = 0.451 | 0.653 |
|  |  | High HC vs Ctrl (d5) | 151 | *t* = −0.489 | 0.626 |
|  |  | Low HC vs Ctrl (d8) | 151 | *t* = 0.304 | 0.761 |
|  |  | High HC vs Ctrl (d8) | 151 | *t* = −0.591 | 0.556 |
| **Exp.** | **Dependent variable** | **Analysis / Effect** | **df / n** | **Statistic** | ***P*** |
| **Exp. 1 – Confirmatory cohort** | Anxiety index | Wilcoxon rank-sum test – low HC | *n* = 30/group | *W* = 333.0 | **0.042** |
|  | Mean velocity |  | *n* = 30/group | *W* = 345.0 | 0.94 |
|  | Shallow arm latency |  | *n* = 30/group | *W* = 382.0 | 0.159 |
|  | Shallow arm frequency |  | *n* = 30/group | *W* = 563.5 | **0.046** |
|  | Shallow arm time |  | *n* = 30/group | *W* = 527.5 | 0.127 |

**Supplementary Table 2.**
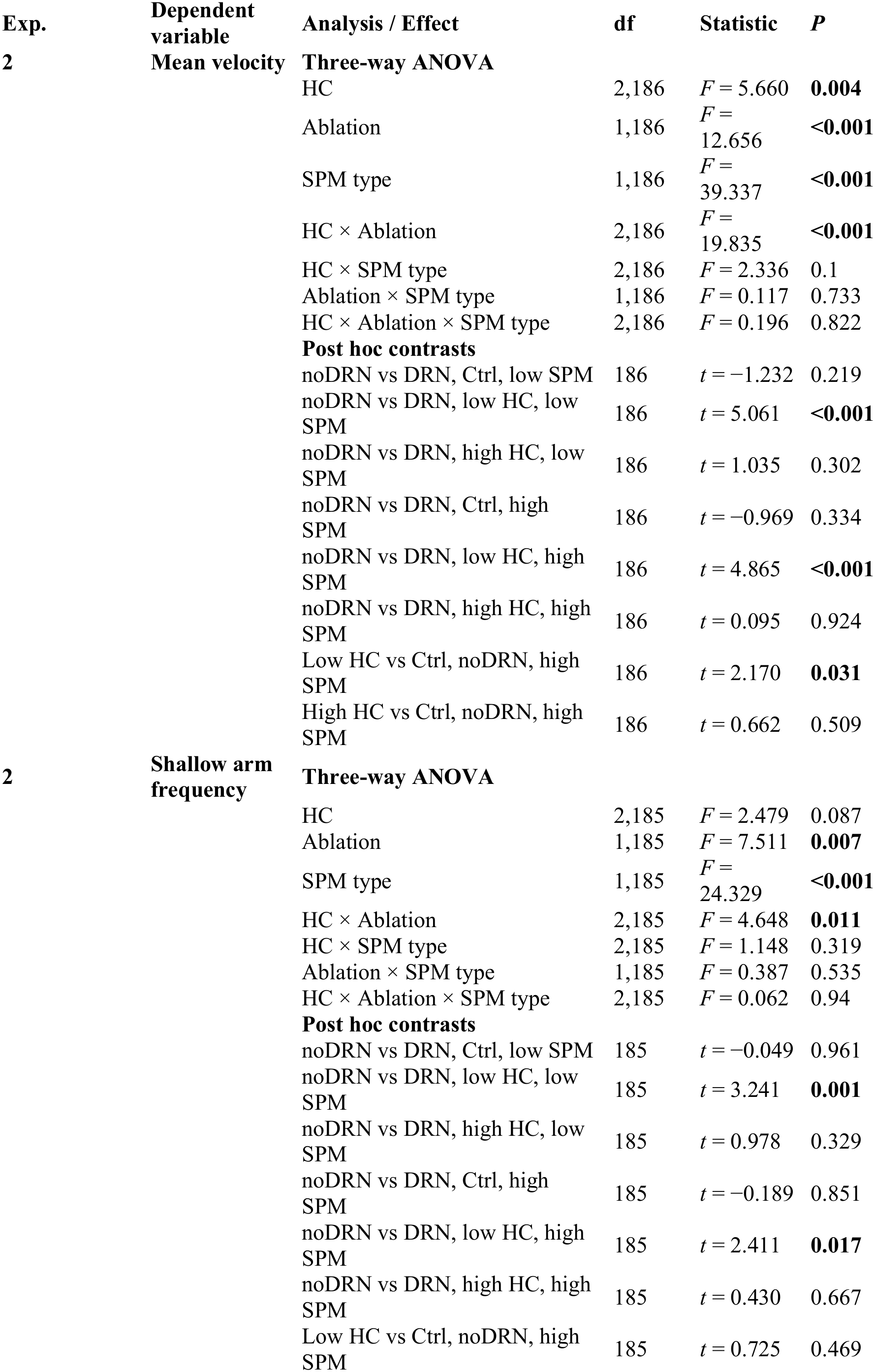
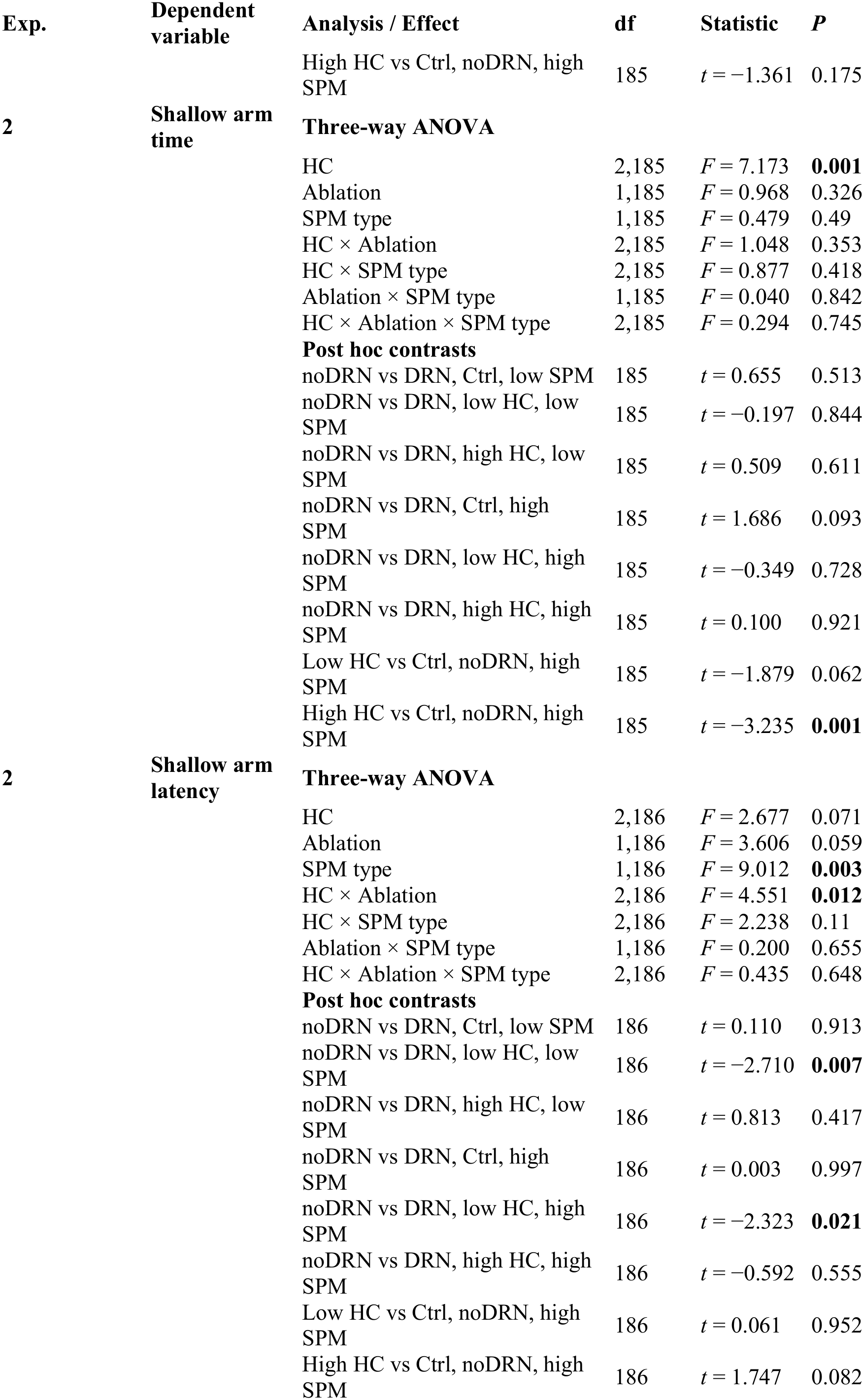
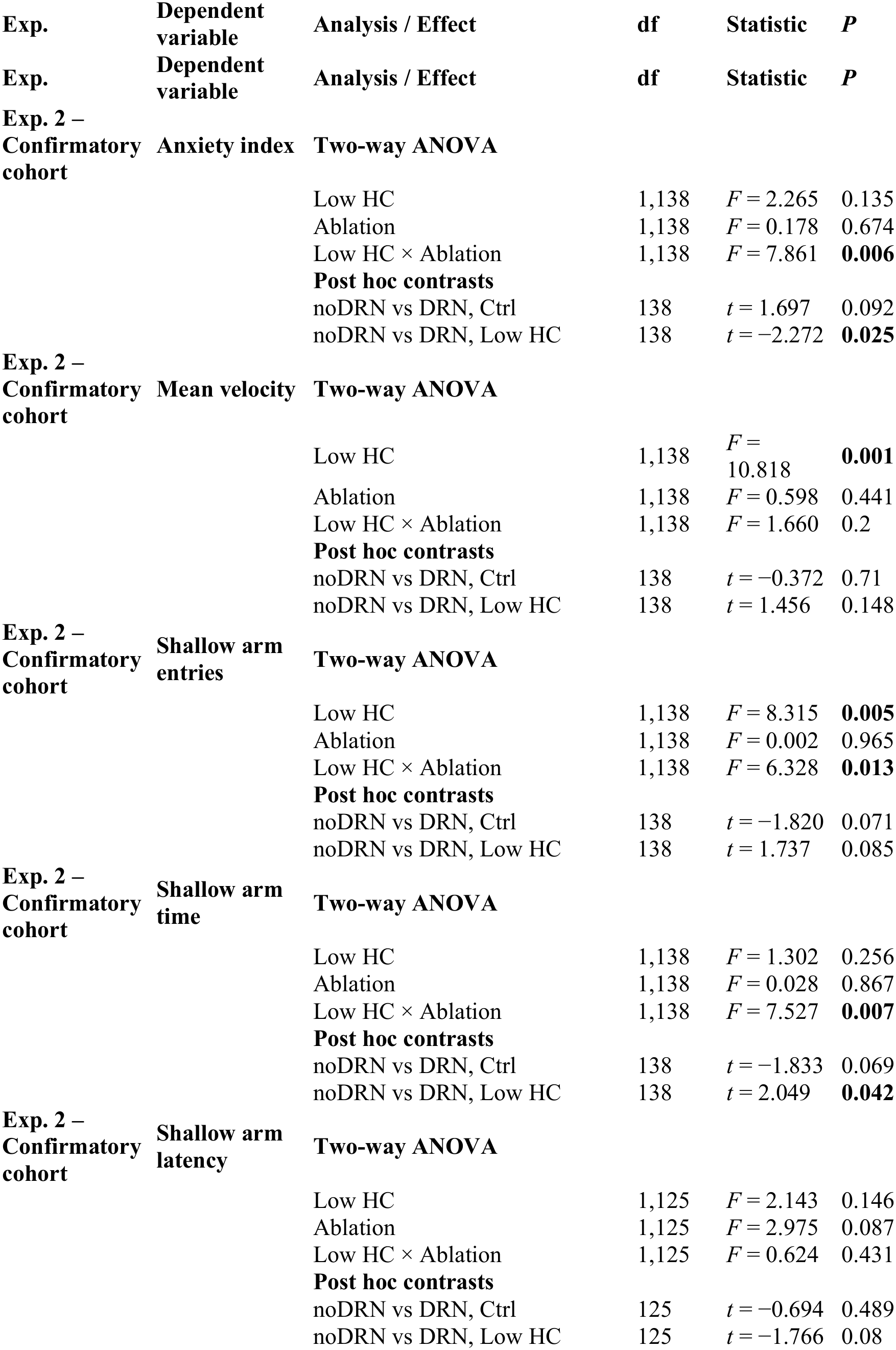

| **Exp.** | **Dependent variable** | **Analysis / Effect** | **df** | **Statistic** | ***P*** |
|---|---|---|---|---|---|
| **2** | **Mean velocity** | **Three-way ANOVA** |  |  |  |
|  |  | HC | 2,186 | *F* = 5.660 | **0.004** |
|  |  | Ablation | 1,186 | *F* = 12.656 | **<0.001** |
|  |  | SPM type | 1,186 | *F* = 39.337 | **<0.001** |
|  |  | HC × Ablation | 2,186 | *F* = 19.835 | **<0.001** |
|  |  | HC × SPM type | 2,186 | *F* = 2.336 | 0.1 |
|  |  | Ablation × SPM type | 1,186 | *F* = 0.117 | 0.733 |
|  |  | HC × Ablation × SPM type | 2,186 | *F* = 0.196 | 0.822 |
|  |  | **Post hoc contrasts** |  |  |  |
|  |  | noDRN vs DRN, Ctrl, low SPM | 186 | *t* = −1.232 | 0.219 |
|  |  | noDRN vs DRN, low HC, low SPM | 186 | *t* = 5.061 | **<0.001** |
|  |  | noDRN vs DRN, high HC, low SPM | 186 | *t* = 1.035 | 0.302 |
|  |  | noDRN vs DRN, Ctrl, high SPM | 186 | *t* = −0.969 | 0.334 |
|  |  | noDRN vs DRN, low HC, high SPM | 186 | *t* = 4.865 | **<0.001** |
|  |  | noDRN vs DRN, high HC, high SPM | 186 | *t* = 0.095 | 0.924 |
|  |  | Low HC vs Ctrl, noDRN, high SPM | 186 | *t* = 2.170 | **0.031** |
|  |  | High HC vs Ctrl, noDRN, high SPM | 186 | *t* = 0.662 | 0.509 |
| **2** | **Shallow arm frequency** | **Three-way ANOVA** |  |  |  |
|  |  | HC | 2,185 | *F* = 2.479 | 0.087 |
|  |  | Ablation | 1,185 | *F* = 7.511 | **0.007** |
|  |  | SPM type | 1,185 | *F* = 24.329 | **<0.001** |
|  |  | HC × Ablation | 2,185 | *F* = 4.648 | **0.011** |
|  |  | HC × SPM type | 2,185 | *F* = 1.148 | 0.319 |
|  |  | Ablation × SPM type | 1,185 | *F* = 0.387 | 0.535 |
|  |  | HC × Ablation × SPM type | 2,185 | *F* = 0.062 | 0.94 |
|  |  | **Post hoc contrasts** |  |  |  |
|  |  | noDRN vs DRN, Ctrl, low SPM | 185 | *t* = −0.049 | 0.961 |
|  |  | noDRN vs DRN, low HC, low SPM | 185 | *t* = 3.241 | **0.001** |
|  |  | noDRN vs DRN, high HC, low SPM | 185 | *t* = 0.978 | 0.329 |
|  |  | noDRN vs DRN, Ctrl, high SPM | 185 | *t* = −0.189 | 0.851 |
|  |  | noDRN vs DRN, low HC, high SPM | 185 | *t* = 2.411 | **0.017** |
|  |  | noDRN vs DRN, high HC, high SPM | 185 | *t* = 0.430 | 0.667 |
|  |  | Low HC vs Ctrl, noDRN, high SPM | 185 | *t* = 0.725 | 0.469 |
| **Exp.** | **Dependent variable** | **Analysis / Effect** | **df** | **Statistic** | ***P*** |
|  |  | High HC vs Ctrl, noDRN, high SPM | 185 | *t* = −1.361 | 0.175 |
| **2** | **Shallow arm time** | **Three-way ANOVA** |  |  |  |
|  |  | HC | 2,185 | *F* = 7.173 | **0.001** |
|  |  | Ablation | 1,185 | *F* = 0.968 | 0.326 |
|  |  | SPM type | 1,185 | *F* = 0.479 | 0.49 |
|  |  | HC × Ablation | 2,185 | *F* = 1.048 | 0.353 |
|  |  | HC × SPM type | 2,185 | *F* = 0.877 | 0.418 |
|  |  | Ablation × SPM type | 1,185 | *F* = 0.040 | 0.842 |
|  |  | HC × Ablation × SPM type | 2,185 | *F* = 0.294 | 0.745 |
|  |  | **Post hoc contrasts** |  |  |  |
|  |  | noDRN vs DRN, Ctrl, low SPM | 185 | *t* = 0.655 | 0.513 |
|  |  | noDRN vs DRN, low HC, low SPM | 185 | *t* = −0.197 | 0.844 |
|  |  | noDRN vs DRN, high HC, low SPM | 185 | *t* = 0.509 | 0.611 |
|  |  | noDRN vs DRN, Ctrl, high SPM | 185 | *t* = 1.686 | 0.093 |
|  |  | noDRN vs DRN, low HC, high SPM | 185 | *t* = −0.349 | 0.728 |
|  |  | noDRN vs DRN, high HC, high SPM | 185 | *t* = 0.100 | 0.921 |
|  |  | Low HC vs Ctrl, noDRN, high SPM | 185 | *t* = −1.879 | 0.062 |
|  |  | High HC vs Ctrl, noDRN, high SPM | 185 | *t* = −3.235 | **0.001** |
| **2** | **Shallow arm latency** | **Three-way ANOVA** |  |  |  |
|  |  | HC | 2,186 | *F* = 2.677 | 0.071 |
|  |  | Ablation | 1,186 | *F* = 3.606 | 0.059 |
|  |  | SPM type | 1,186 | *F* = 9.012 | **0.003** |
|  |  | HC × Ablation | 2,186 | *F* = 4.551 | **0.012** |
|  |  | HC × SPM type | 2,186 | *F* = 2.238 | 0.11 |
|  |  | Ablation × SPM type | 1,186 | *F* = 0.200 | 0.655 |
|  |  | HC × Ablation × SPM type | 2,186 | *F* = 0.435 | 0.648 |
|  |  | **Post hoc contrasts** |  |  |  |
|  |  | noDRN vs DRN, Ctrl, low SPM | 186 | *t* = 0.110 | 0.913 |
|  |  | noDRN vs DRN, low HC, low SPM | 186 | *t* = −2.710 | **0.007** |
|  |  | noDRN vs DRN, high HC, low SPM | 186 | *t* = 0.813 | 0.417 |
|  |  | noDRN vs DRN, Ctrl, high SPM | 186 | *t* = 0.003 | 0.997 |
|  |  | noDRN vs DRN, low HC, high SPM | 186 | *t* = −2.323 | **0.021** |
|  |  | noDRN vs DRN, high HC, high SPM | 186 | *t* = −0.592 | 0.555 |
|  |  | Low HC vs Ctrl, noDRN, high SPM | 186 | *t* = 0.061 | 0.952 |
|  |  | High HC vs Ctrl, noDRN, high SPM | 186 | *t* = 1.747 | 0.082 |
| **Exp.** | **Dependent variable** | **Analysis / Effect** | **df** | **Statistic** | ***P*** |
| **Exp.** | **Dependent variable** | **Analysis / Effect** | **df** | **Statistic** | ***P*** |
| **Exp. 2 – Confirmatory cohort** | **Anxiety index** | **Two-way ANOVA** |  |  |  |
|  |  | Low HC | 1,138 | *F* = 2.265 | 0.135 |
|  |  | Ablation | 1,138 | *F* = 0.178 | 0.674 |
|  |  | Low HC × Ablation | 1,138 | *F* = 7.861 | **0.006** |
|  |  | **Post hoc contrasts** |  |  |  |
|  |  | noDRN vs DRN, Ctrl | 138 | *t* = 1.697 | 0.092 |
|  |  | noDRN vs DRN, Low HC | 138 | *t* = −2.272 | **0.025** |
| **Exp. 2 – Confirmatory cohort** | **Mean velocity** | **Two-way ANOVA** |  |  |  |
|  |  | Low HC | 1,138 | *F* = 10.818 | **0.001** |
|  |  | Ablation | 1,138 | *F* = 0.598 | 0.441 |
|  |  | Low HC × Ablation | 1,138 | *F* = 1.660 | 0.2 |
|  |  | **Post hoc contrasts** |  |  |  |
|  |  | noDRN vs DRN, Ctrl | 138 | *t* = −0.372 | 0.71 |
|  |  | noDRN vs DRN, Low HC | 138 | *t* = 1.456 | 0.148 |
| **Exp. 2 – Confirmatory cohort** | **Shallow arm entries** | **Two-way ANOVA** |  |  |  |
|  |  | Low HC | 1,138 | *F* = 8.315 | **0.005** |
|  |  | Ablation | 1,138 | *F* = 0.002 | 0.965 |
|  |  | Low HC × Ablation | 1,138 | *F* = 6.328 | **0.013** |
|  |  | **Post hoc contrasts** |  |  |  |
|  |  | noDRN vs DRN, Ctrl | 138 | *t* = −1.820 | 0.071 |
|  |  | noDRN vs DRN, Low HC | 138 | *t* = 1.737 | 0.085 |
| **Exp. 2 – Confirmatory cohort** | **Shallow arm time** | **Two-way ANOVA** |  |  |  |
|  |  | Low HC | 1,138 | *F* = 1.302 | 0.256 |
|  |  | Ablation | 1,138 | *F* = 0.028 | 0.867 |
|  |  | Low HC × Ablation | 1,138 | *F* = 7.527 | **0.007** |
|  |  | **Post hoc contrasts** |  |  |  |
|  |  | noDRN vs DRN, Ctrl | 138 | *t* = −1.833 | 0.069 |
|  |  | noDRN vs DRN, Low HC | 138 | *t* = 2.049 | **0.042** |
| **Exp. 2 – Confirmatory cohort** | **Shallow arm latency** | **Two-way ANOVA** |  |  |  |
|  |  | Low HC | 1,125 | *F* = 2.143 | 0.146 |
|  |  | Ablation | 1,125 | *F* = 2.975 | 0.087 |
|  |  | Low HC × Ablation | 1,125 | *F* = 0.624 | 0.431 |
|  |  | **Post hoc contrasts** |  |  |  |
|  |  | noDRN vs DRN, Ctrl | 125 | *t* = −0.694 | 0.489 |
|  |  | noDRN vs DRN, Low HC | 125 | *t* = −1.766 | 0.08 |

**Supplementary table 3.**

| **Exp.** | **Dependent variable** | **Analysis / Effect** | **df** | **Statistic** | ***P*** |
|---|---|---|---|---|---|
| **Exp. 3** | **Anxiety index** | **Three-way ANOVA** |  |  |  |
|  |  | Low HC | 1,103 | *F* = 1.145 | 0.287 |
|  |  | Ablation | 1,103 | *F* = 1.437 | 0.233 |
|  |  | Dopamine antagonist | 2,103 | *F* = 1.809 | 0.169 |
|  |  | Ablation × Dopamine antagonist | 2,103 | *F* = 3.178 | **0.046** |
|  |  | **Post hoc contrasts** |  |  |  |
|  |  | D1 vs Vehicle, DRN | 103 | *t* = −0.529 | 0.598 |
|  |  | D2 vs Vehicle, DRN | 103 | *t* = −0.481 | 0.631 |
|  |  | D1 vs Vehicle, noDRN | 103 | *t* = 3.017 | **0.003** |
|  |  | D2 vs Vehicle, noDRN | 103 | *t* = 0.579 | 0.564 |
| **Exp. 3** | **Mean velocity** | **Three-way ANOVA** |  |  |  |
|  |  | Low HC | 1,103 | *F* = 1.225 | 0.271 |
|  |  | Ablation | 1,103 | *F* = 6.463 | **0.013** |
|  |  | Dopamine antagonist | 2,103 | *F* = 14.601 | **<0.001** |
|  |  | Ablation × Dopamine antagonist | 2,103 | *F* = 0.173 | 0.842 |
|  |  | **Post hoc contrasts** |  |  |  |
|  |  | D1 vs Vehicle, DRN | 103 | *t* = −3.823 | **<0.001** |
|  |  | D2 vs Vehicle, DRN | 103 | *t* = 0.101 | 0.92 |
|  |  | D1 vs Vehicle, noDRN | 103 | *t* = −3.019 | **0.003** |
|  |  | D2 vs Vehicle, noDRN | 103 | *t* = 0.303 | 0.762 |
| **Exp. 3** | **Shallow arm entries** | **Three-way ANOVA** |  |  |  |
|  |  | Low HC | 1,103 | *F* = 0.878 | 0.351 |
|  |  | Ablation | 1,103 | *F* = 3.738 | 0.056 |
|  |  | Dopamine antagonist | 2,103 | *F* = 4.529 | **0.013** |
|  |  | Ablation × Dopamine antagonist | 2,103 | *F* = 0.234 | 0.792 |
|  |  | **Post hoc contrasts** |  |  |  |
|  |  | D1 vs Vehicle, DRN | 103 | *t* = −1.376 | 0.172 |
|  |  | D2 vs Vehicle, DRN | 103 | *t* = 0.376 | 0.708 |
|  |  | D1 vs Vehicle, noDRN | 103 | *t* = −2.347 | **0.021** |
|  |  | D2 vs Vehicle, noDRN | 103 | *t* = 0.022 | 0.983 |
| **Exp. 3** | **Shallow arm time** | **Three-way ANOVA** |  |  |  |
|  |  | Low HC | 1,103 | *F* = 0.048 | 0.827 |
|  |  | Ablation | 1,103 | *F* = 1.200 | 0.276 |
|  |  | Dopamine antagonist | 2,103 | *F* = 1.575 | 0.212 |
|  |  | Ablation × Dopamine antagonist | 2,103 | *F* = 4.603 | **0.012** |
|  |  | **Post hoc contrasts** |  |  |  |
|  |  | D1 vs Vehicle, DRN | 103 | *t* = 3.344 | **0.001** |
|  |  | D2 vs Vehicle, DRN | 103 | *t* = 1.402 | 0.164 |
|  |  | D1 vs Vehicle, noDRN | 103 | *t* = −0.876 | 0.383 |
|  |  | D2 vs Vehicle, noDRN | 103 | *t* = −0.929 | 0.355 |
| **Exp. 3** | **Shallow arm latency** | **Three-way ANOVA** |  |  |  |
|  |  | Low HC | 1,93 | *F* = 1.492 | 0.225 |
|  |  | Ablation | 1,93 | *F* = 5.559 | **0.02** |
|  |  | Dopamine antagonist | 2,93 | *F* = 1.220 | 0.3 |
|  |  | Ablation × Dopamine antagonist | 2,93 | *F* = 3.308 | **0.041** |
|  |  | **Post hoc contrasts** |  |  |  |
|  |  | D1 vs Vehicle, DRN | 93 | *t* = −0.659 | 0.512 |
|  |  | D2 vs Vehicle, DRN | 93 | *t* = 0.337 | 0.737 |
|  |  | D1 vs Vehicle, noDRN | 93 | *t* = 2.835 | **0.006** |
|  |  | D2 vs Vehicle, noDRN | 93 | *t* = 0.613 | 0.542 |

